# Optogenetic Proximity Labeling Maps Spatially Resolved Mitochondrial Surface Proteomes and a Locally Regulated Ribosome Pool

**DOI:** 10.64898/2025.12.21.693523

**Authors:** Chulhwan S. Kwak, Zehui Du, Joseph S. Creery, Emily M. Wilkerson, Michael B. Major, Joshua E. Elias, Xinnan Wang

**Affiliations:** Department of Neurosurgery, Stanford University School of Medicine, Stanford, CA, USA; Biohub, San Francisco, CA, USA; Department of Cell Biology and Physiology, Department of Otolaryngology, Washington University in St. Louis, St. Louis, MO, USA

## Abstract

Outer mitochondrial membranes (OMM) function as dynamic hubs for inter-organelle communication, integrating bidirectional signals, and coordinating organelle behavior in a context-dependent manner. However, tools for mapping mitochondrial surface proteomes with high spatial and temporal resolution remain limited. Here, we introduce an optogenetic proximity labeling strategy using LOV-Turbo, a light-activated biotin ligase, to profile mitochondrial surface proteomes with improved precision, temporal control, and reduced background. By fusing LOV-Turbo to a panel of variants of an OMM-anchored protein, Miro1, we generate spatially distinct baits that resolve modular architectures and regulatory states of the OMM proteomes across diverse conditions, a database we name MitoSurf. Building on this proteomic map, we present RiboLOOM, a platform that defines LOV-Turbo labeled ribosomes and their bound mRNAs at the mitochondrial surface. MitoSurf and RiboLOOM uncover a spatially distinct ribosome pool at the OMM that is maintained by Miro1, enabling local mRNA engagement and translation of mitochondria-related proteins. These findings establish Miro1 as a key organizer of mitochondrial protein biogenesis through spatial confinement of surface-associated ribosomes. Our platform reveals an uncharted layer of mitochondrial surface biology and provides a generalizable strategy to dissect dynamic RNA-protein-organelle interfaces in living cells.

## Introduction

Mitochondria are highly mobile organelles, zipping around and shapeshifting constantly within the cell ^1,2^. Beyond their classical role in ATP production, mitochondria serve as signaling hubs ^3^, and their outer membrane acts as a key interface for communication with other organelles and the extracellular milieu. The mitochondrial surface is equipped with membrane channels that regulate the exchange of proteins, ions, and metabolites, as well as tethering complexes that physically link mitochondria to the endoplasmic reticulum (ER), vesicles, and cytoskeletal tracks. These molecular interactions are governed by a specialized mitochondrial surface proteome that enables cell-type-specific mitochondrial behaviors, homeostasis, and adaptive responses. Although techniques such as proximity labeling-mass spectrometry (PL-MS), affinity purification-mass spectrometry (AP-MS), and organelle-specific interactome mapping have provided valuable insights into mitochondrial proteomes and contact sites ^4–10^, the spatial resolution of discrete protein modules on the outer mitochondrial membrane (OMM) remains limited. Moreover, how the mitochondrial surface proteome dynamically responds to genetic or pharmacological perturbations in a cell-type-specific manner is poorly understood. Addressing these questions is critical to uncovering the mechanisms of mitochondrial plasticity and inter-organelle communication.

Proximity labeling (PL) has emerged as a powerful approach for defining the local proteome surrounding a target protein in living cells, complementing affinity-based proteomic strategies ^11,12^. Among PL systems, TurboID, an engineered biotin ligase with fast kinetics and constitutive activity, has enabled robust proteome labeling across diverse biological systems ^12^. However, its high background labeling in certain cell types, such as neurons, poses a challenge due to elevated levels of endogenous biotin, often present in biotin-supplemented media essential for neuronal survival ^13,14^. APEX, a peroxidase-based PL system, requires hydrogen peroxide to initiate biotinylation, which may induce oxidative stress and cytotoxicity, particularly in neurons or cancer cells, both of which are intrinsically susceptible to redox imbalance ^11,15,16^. BmTyr ^17^ and LaccID ^18^ offer hydrogen peroxide-free PL, but both are currently limited to cell surface proteomics due to their reliance on membrane-impermeable phenol probes and lack of compatibility with intracellular environments. To overcome these limitations, a recently developed light-gated variant of TurboID, known as LOV-Turbo (LOV-Tb), enables precise temporal control of PL at intracellular organelles ^19^. In the absence of light, LOV-Tb remains inactive, initiating biotinylation only upon blue light stimulation. This temporal gating is particularly advantageous for capturing proteomic changes triggered by rapid and transient stimuli, such as neuronal activation or acute toxic insults. By precisely timing biotinylation events, LOV-Tb allows selective labeling of proteomic states before or after defined perturbations, thereby minimizing background and enhancing specificity in cell-type- and context-specific proteomic profiling.

Another major limitation in defining the mitochondrial surface proteome is the lack of an accurate and comprehensive annotation database. Existing resources such as MitoCarta3.0 ^7^ and native organelle immunoprecipitation (IP)-based proteomes ^5,10^ serve as valuable tools for researchers seeking mitochondrial protein references across diverse biological contexts. However, these databases have notable gaps. MitoCarta3.0, while widely used, primarily catalogs proteins localized to the mitochondrial matrix or inner membrane, with only partial inclusion of proteins on the OMM. Moreover, it lacks submitochondrial resolution, making it difficult to distinguish proteins truly residing at the organelle surface from those embedded deeper within. The native organelle IP approach offers broader coverage, including proteomic data for multiple subcellular compartments such as mitochondria, lysosomes, nucleus, cytosol, P-bodies, and stress granules. Nevertheless, this method does not provide a curated, surface-specific mitochondrial proteome. To fully understand how the mitochondrial surface acts as a dynamic interface for inter-organelle signaling and cellular adaptation, there is a pressing need for high-resolution, surface-specific mitochondrial proteomic maps. Establishing such a resource will be essential not only for uncovering the fundamental biology of organelle communication but also for guiding therapeutic strategies.

To investigate cell-type-specific signatures of the mitochondrial surface proteome, we selected the OMM protein Miro1 as the scaffold for constructing optogenetic proximity labeling baits, due to its unique topology and multifunctional roles. Miro1 is anchored to the OMM via a C-terminal transmembrane (TM) domain and pieces into the intermembrane space (IMS) via a short segment (C-tail), which interacts with the mitochondrial contact site and cristae organizing system (MICOS) complex on the inner mitochondrial membrane (IMM) ^20,21^. Its N-terminal region, exposed to the cytosol, contains two calcium-binding motifs and two GTPase domains. This modular architecture enables Miro1 to serve both as a mitochondrial receptor for facilitating docking of mitochondria to the ER and cytoskeletal motors, and as a regulatory switch that integrates intracellular calcium and GTP signals to modulate mitochondrial behavior. Miro1 is also a key regulator of mitophagy, the selective autophagic degradation of mitochondria. Studies have shown that removal of Miro1 from the OMM marks, segregates, and quarantines dysfunctional mitochondria destined for mitophagy ^22–25^. To dissect the molecular organization of the OMM in different cell contexts, we engineered a series of LOV-Tb-conjugated Miro1 variants, each targeting distinct subdomains or functional states of the protein (Figure 1A). Using this panel, we aimed to resolve sub-compartments of the OMM and characterize mitochondrial surface proteome plasticity in a cell-type- and perturbation-dependent manner. We applied this system across four distinct human cell lines: glioblastoma (GBM) cells, induced pluripotent stem cell (iPSC)-induced neurons (iN) (healthy control and isogenic *SNCA-A53T*, a prominent Parkinson’s disease variant), as well as HEK293T cells, a widely used standard for proteomics benchmarking (Figure 1B) ^5^. Leveraging this platform, we uncover a spatially resolved and dynamic mitochondrial surface proteome, reveal its plasticity across cell types and perturbations, and identify previously unrecognized functions of Miro1 in regulating mitochondrial interface biology.

**Figure 1.**
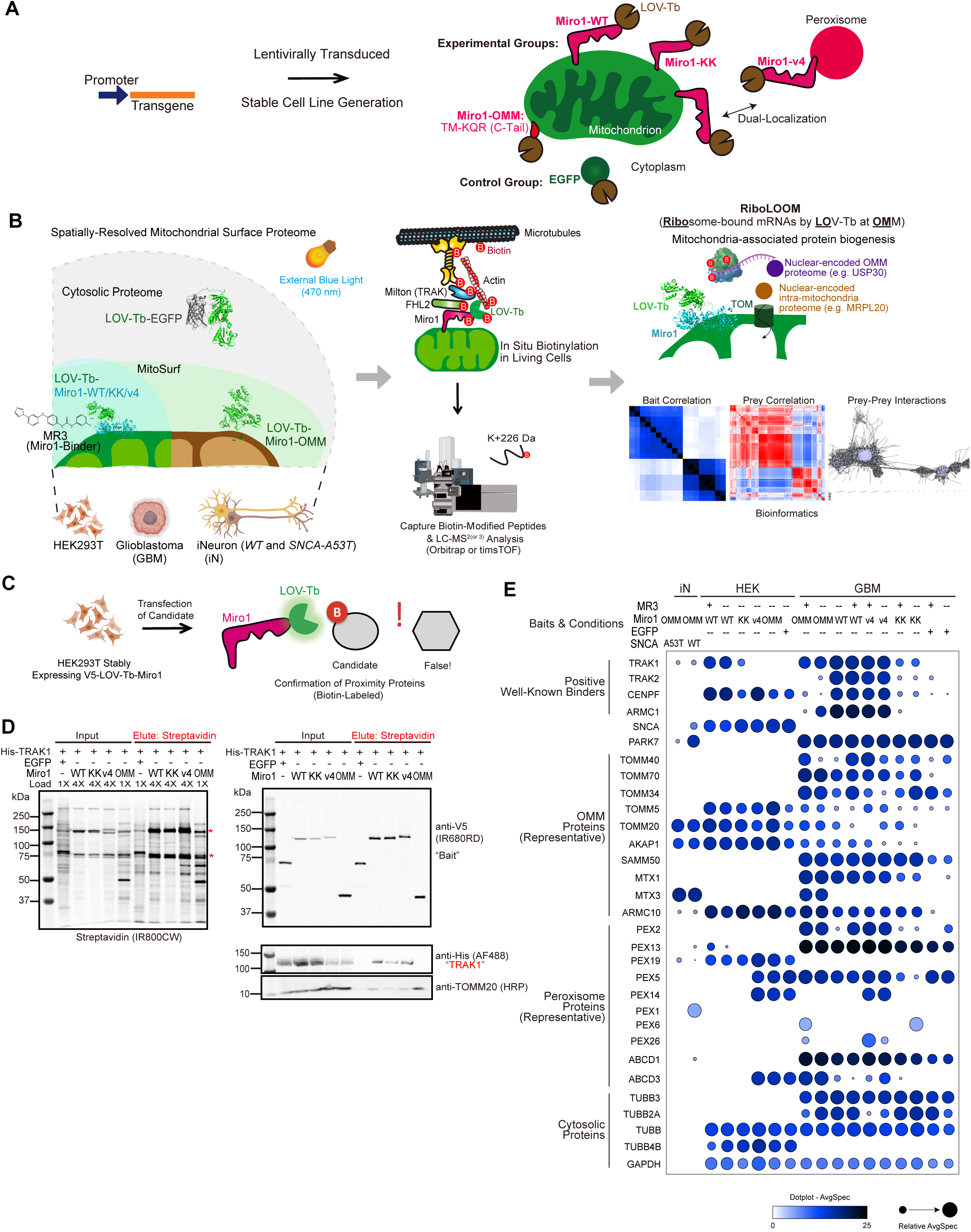
Workflow and validation of our optogenetic proximity labeling method. (A) Schematic representation of construct and stable cell line generation. (B) Overall workflow. Upon light irradiation (470 nm), proximal proteins near the baits are biotinylated, digested, and captured for mass spectrometry analysis to identify biotin-modified peptides. Proteomic hits are further analyzed by bioinformatics and predictive structural methods. Simultaneously, biotin-labeled ribosomes are purified, and their bound mRNAs are analyzed by RT-qPCR. (C) A cartoon to depict the in situ biotinylation assay using the optogenetic proximity labeling toolkit (LOV-Tb) in living HEK293T cells. (D) The in situ biotinylation assay on well-known interacting partners (TRAK1 and TOMM20) of Miro1 by the platform as shown in (C). Biotinylated protein pools were eluted from Streptavidin beads. * indicates endogenous biotinylated proteins. (E) Normalized mean MS intensity of selected proteins identified across baits and cell types is shown in dot plot. Color density shows intensity, and circle size denotes relative abundance (AvgSpec for a given bait divided by the maximal AvgSpec recorded for that prey across all baits, scaled 0-1).

## Result

### Optogenetic proximity labeling profiles mitochondrial surface proteomes across cell types

Our panel of Miro1-based baits included: (1) Miro1-WT, the full-length wild-type protein; (2) Miro1-KK, an EF-hand mutant deficient in calcium binding; (3) Miro1-OMM, a truncated variant containing only the C-terminal TM domain and the IMS-facing C-tail (KQR motif); and (4) Miro1-v4, a peroxisome-localized isoform previously reported to co-target both mitochondria and peroxisomes ^26, 27^ (Figure 1A, S1A, S2, Table S1). We used EGFP as a cytosolic control. In selected conditions, we also applied Miro1 Reducer 3 (MR3), a small molecule that binds the C-terminal GTPase domain of Miro1, to assess its impact on mitochondrial surface protein composition ^20,28,29^ (Figure 1B). Co-immunostaining and live imaging confirmed that all Miro1 constructs localized correctly to mitochondria or peroxisomes, and did not affect their morphology (Figure S1A, S2) or the mitochondrial membrane potential detected by TMRM normalized to MitoTracker Green (Figure S1B). Following blue light activation, in situ biotinylation was performed in living cells. Both immunostaining and streptavidin pull-down followed by Western blotting validated efficient biotinylation across all LOV-Tb-tagged baits, indicating robust labeling activity (Figure S1C-S2).

Streptavidin-enriched proteins were profiled by LC-MS/MS (Figure 1B, S3A-D). Each bait condition was analyzed in 3-4 biological replicates. Pairwise Pearson correlations across replicates ranged from 0.71 to 0.97, reflecting high reproducibility. Principal component analysis (PCA) further confirmed clear separation of samples based on bait identity and cell type (Figure S3E), underscoring the biological specificity. To complement proximity proteomics, we also performed whole-cell label-free or TMT proteomic profiling using both data-independent (DIA) and data-dependent (DDA) acquisition across key experimental contexts (Figure S4). In total, 93 mass spectrometry samples were processed (Table S2-S3), enabling integrative assessment of mitochondrial surface remodeling at both the spatial and global proteome resolution (all data are publicly available via a user-friendly browser called MiroScape – https://miroscape.github.io/MiroScape/#/home) ^30^.

To validate the proximity labeling results, we performed targeted Western blot analysis of two well-established Miro1-associated proteins: TRAK1 and TOMM20. Their relative abundance across LOV-Tb-Miro1 baits (Figure 1C-D) closely mirrored the patterns observed in our mass spectrometry data (Figure 1E), supporting the reliability of our PL method. In addition to these, multiple known OMM components, including TOMM40, SAMM50, and TOMM70, as well as other Miro1 interactors such as CENPF ^31^ TRAK2 ^32^, and ARMC1 ^33^, were specifically enriched (Figure 1E).

Using this platform, we identified 1,838 proteins aggregated across all Miro1-derived baits and cell types (Figure 2A), representing OMM resident proteins and surface-proximal interactors during transient contact or import. We named our expanded proteome **MitoSurf** (**<u>Mito</u>**chondrial **<u>Surf</u>**ace Proteome; publicly available on MiroScape – https://miroscape.github.io/MiroScape/#/mitoSurf) ^30^. Among **MitoSurf** preys, we calculated a High-Confidence Proximity Score (HCPS), accounting for factors including global background, statistical significance, replicate consistency, enrichment relative to EGFP, and subcellular localization from known annotation databases (Figure 2A, Table S2, Methods). The values of HCPS correlated with the confidence of OMM resident proteins or preys enriched at the OMM. Cross-referencing with MitoCarta3.0 revealed that only 183 of **MitoSurf** were annotated as mitochondrial proteins and a substantially larger set of additional proteins were not present in MitoCarta3.0 (Figure 2A), suggesting enrichment of previously uncharacterized, multi-organelle, or interface-associated components. Sub-organelle analysis showed that 50 were annotated as OMM proteins, while others mapped to the matrix, IMM, and IMS, suggesting that our approach may also capture the transport of mitochondrially targeted peptides at or near the OMM (details later) (Figure 2A). Comparison with native organelle IP datasets confirmed the inclusion of high-confidence mitochondrial and multi-compartmental proximity interactors (Figure 2A), validating the spatial breadth and specificity of our dataset. Overall, a substantial number of identified proteins in **MitoSurf** were not annotated in either MitoCarta3.0 or the organelle IP reference datasets, including established Miro1-interacting partners such as TRAK1 and TRAK2, highlighting limitations in current sub-organelle annotation resources. Hence, **MitoSurf** reflects the unique capability of LOV-Tb-Miro1 labeling to resolve a broader and more dynamic landscape of mitochondrial surface interactions and serves as a new resource for the community.

**Figure 2.**
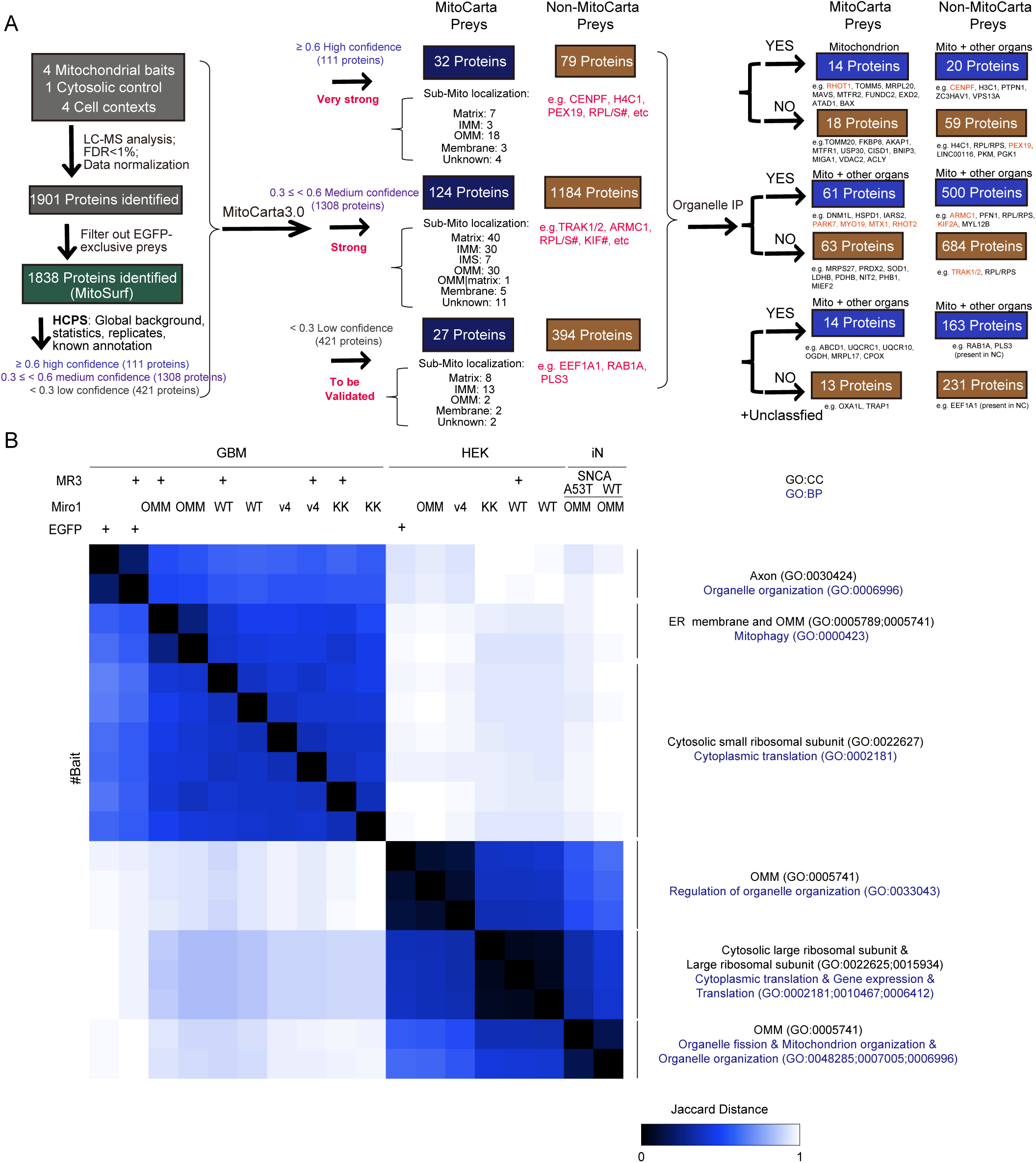
A comprehensive mitochondrial surface proteome by LOV-Tb and bait self-organization. (A) Summary of our overall dataset and quality controls. The number of proteins and representative proteins annotated with MitoCarta3.0 ^2^ and organelle IP ^3^ are shown. (B) Bait-Bait similarity heatmap displays the Jaccard distance score (0-1) and hierarchical clustering. Bait is labeled on the top of the heatmap. Each bait was manually profiled in g:profiler and the identified GO (BP: Biological Process) and GO (CC: Cellular Compartment) terms are indicated to the right of the heatmap.

To assess the similarity of proximity labeling patterns across different baits and cell types, we calculated Jaccard distances based on the overlap of preys by each bait, followed by hierarchical clustering (Figure 2B). In this analysis, a lower Jaccard distance indicates a greater degree of prey overlap between baits, reflecting more similar labeling patterns. Accordingly, baits that localize to similar subcellular regions or shared structural features cluster together, whereas functionally or spatially distinct baits exhibit greater distance. In GBM cells, all Miro1-derived baits clustered distinctly from the cytosolic EGFP control, whereas in HEK293T cells, Miro1-OMM and Miro1-v4 baits showed greater overlap with EGFP, suggesting partial convergence between cytosolic and mitochondrial surface labeling in this context. These results highlight both structural and context-dependent plasticity in mitochondrial surface proteome composition. Notably, Miro1-v4 bait was closely clustered with other Miro1 baits (Figure 2B, 3E) and did not label exclusive preys (Table S2), reflecting its mitochondrial localization (Figure S1-S2).

Together, we demonstrate the LOV-Tb-Miro1 bait suite as a robust tool for mapping the mitochondrial surface proteome with high spatial resolution and temporal control. The resulting dataset, **MitoSurf**, establishes a framework for dissecting mitochondrial signaling pathways and organelle interactions that we will explore in the subsequent sections.

### Prey-prey correlation analysis assigns specialized modules

To identify functional modules within the mitochondrial surface proteome, we performed prey-prey correlation analysis across all proximity-labeled proteins ^4^. Hierarchical clustering based on Pearson correlation coefficients grouped preys into distinct clusters, suggesting the presence of co-regulated or spatially co-localized protein assemblies (Figure 3A). Gene Ontology (GO) enrichment analysis of individual clusters revealed diverse and functionally coherent modules. These included well-defined categories such as cytoskeleton organization, organelle membrane assembly, mitochondrial organization, and cytoplasmic translation (Figure 3A).

**Figure 3.**
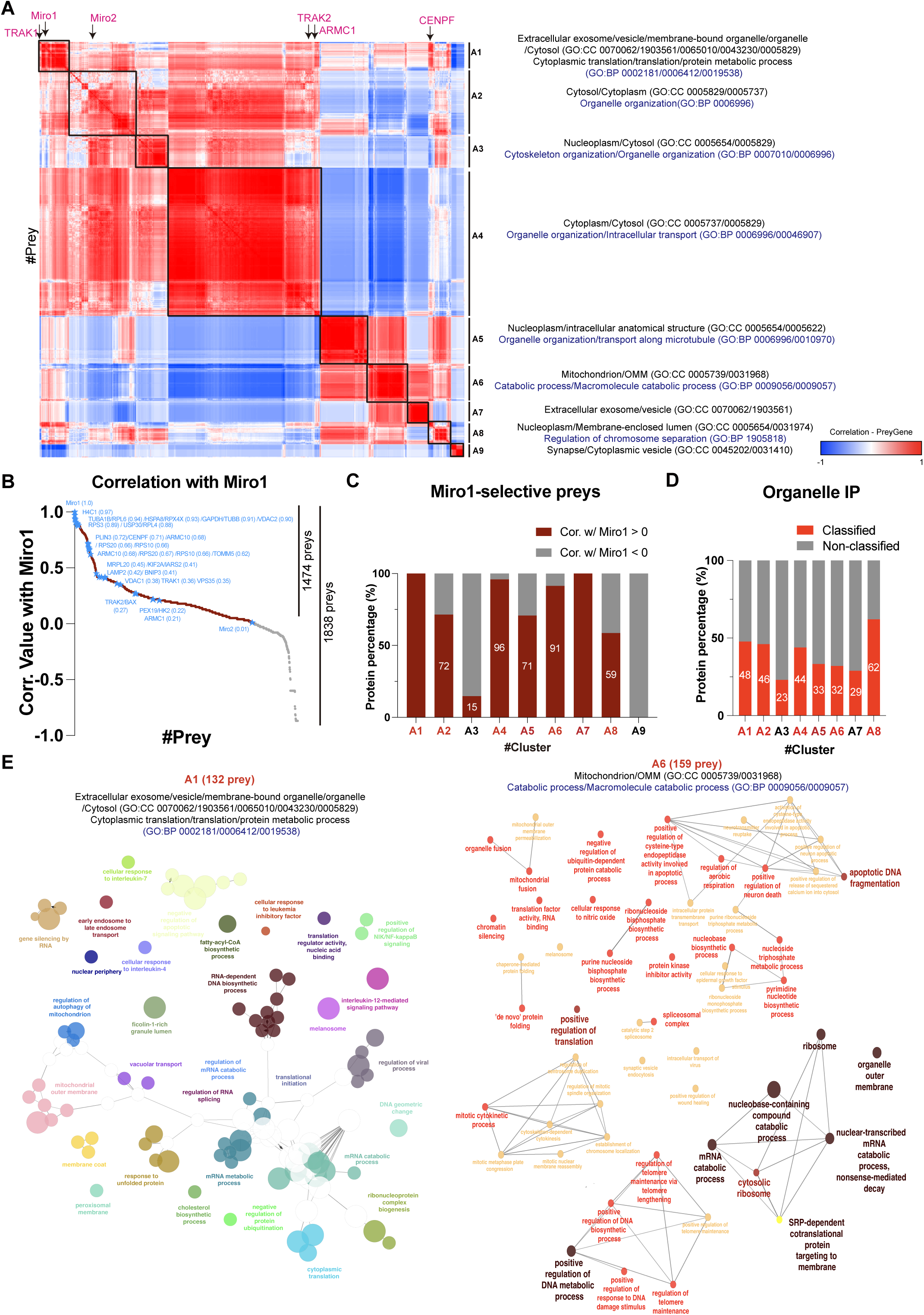
Prey-prey correlation analysis. (A) A heat map of the entire network depending on Pearson’s correlation shows top GO terms in each clustered group listed from g:profiler. Correlation value (each prey genes) is scaled -1 to +1: deep red is +1 (strong positive), white is 0 (no correlation), and deep blue is -1 (strong negative). Six sub-clusters (A1 to A6) are shown with the corresponding GO terms (CC: Cellular component and BP: Biological process). Proteins labeled on the top of the heat map are representative well-known interacting partners of Miro1. (B) The correlation values of all identified preys with Miro1. Self-correlation value (Miro1-Miro1) is 1. Representative hits are highlighted in blue. (C) Bar graph shows the number of Miro1-correlated preys in each distinct cluster. (D) Stacked bar shows the proportion and number of proteins that could be assigned to organelle IP (red). (E) The subcellular localization of proteins in selected clusters (132 proteins in A1 and 159 proteins in A6). The ClueGO analysis (GO: Cellular Component) under Cytoscape plugin shows Miro1-associated preys self-organize into distinct modules. The nodes represent significantly enriched GO terms/pathways. Each color denotes a specific functional group assigned by ClueGO. Node sizes are proportional to the - log_10_ adjusted p values (larger is more significant).

Prey-prey correlation analysis is a powerful strategy not only for distinguishing functional subcellular compartments but also for uncovering putative associations between distinct biological modules ^4^. Since all our LOV-Tb baits were anchored on the Miro1 backbone, we reasoned that a subset of correlated prey proteins would reflect Miro1-dependent biological processes. To investigate this, we assessed the correlation of Miro1, as a prey, with other preys across both on-diagonal and off-diagonal clusters (Figure 3A-B). We identified several functional modules with high (correlation > 0.6) or moderate (correlation > 0.19) association with Miro1 (Figure 3B), including known Miro1-interacting partners such as TRAK1/2 and ARMC1, supporting the specificity and biological relevance of our labeling strategy (Figure 3B-D).

Miro1-associated preys were mapped to multiple organelles and functional modules (Figure 3C-D), corroborating the multi-compartmental nature of Miro1 interactions. Pathway network analysis (ClueGO) of Miro1-correlated preys revealed enrichment in well-established Miro1 functions, such as mitochondrial membrane and organelle transport, but also uncovered unexpected pathways, including extracellular exosomes, ribonucleoprotein complexes, and cytoplasmic translation (Figure 3D). These findings demonstrate the power of our LOV-Tb platform to reveal both known and previously unrecognized biology linked to OMM proteins such as Miro1.

### The mitochondrial surface proteome adapts to cellular environment

Given that all Miro1 baits face the cytosol, we next asked whether the mitochondrial surface proteome is distinguishable from the general cytosolic proteome. We compared proximity-labeled proteins from the cytosolic bait EGFP with those from Miro1-OMM in HEK293T and GBM cells. A partial overlap was observed (Figure 4A, C), suggesting that a fraction of labeled proteins resides at the mitochondria-cytosol interface and can be captured by both probes. However, differential enrichment analysis revealed distinct proteomic signatures: 791 and 368 proteins were significantly enriched in the EGFP bait, while 114 and 338 were enriched in the Miro1-OMM bait, in HEK and GBM cells, respectively (Figure 4A-D). These data indicate that the cytosolic and mitochondrial surface proteomes are at least partially separable in both cell types.

**Figure 4.**
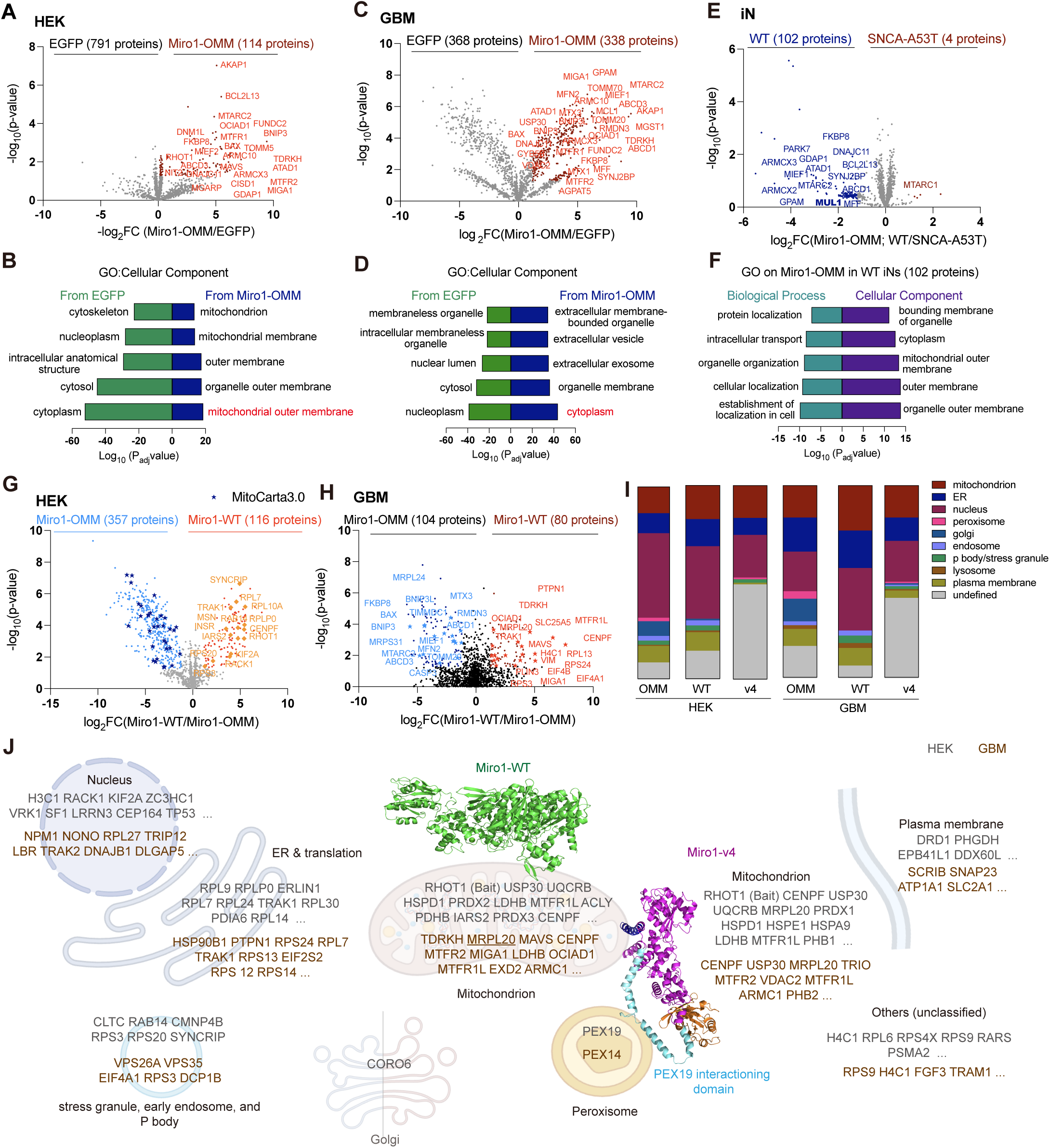
The cell-type-specific proteomes and multi-organellar contacts at the mitochondrial surface. (A, C, E) The volcano plot of prey clusters labeled from Miro1-OMM against those from EGFP in HEK293T (A), GBM (B), and from Miro1-OMM in WT and SNCA-A53T iNs (E). Some proteins annotated by MitoCarta3.0 (OMM or membrane) are highlighted in red in (A, C) or in blue in (E). x axis is log_2_ fold change (FC) between Miro1-OMM and EGFP (right=enriched in Miro1-OMM, left=enriched in EGFP), or between WT and SNCA-A53T. y axis indicates -log_10_ adjusted p value from the statistical test (higher=more significant). Filtration condition is p value<0.05 and log_2_FC≥1.0. Grey-colored dots are not significant. (B, D) The annotated GO terms (Cellular Component) from filtered preys in (A, C). Bars correspond to the GO terms that are significantly enriched in the indicated dataset against the other generated from g:profiler. Bar length is proportional to the enrichment statistics (-log_10_ adjusted p value); longer bars indicate more strongly enriched terms. The bars are sorted in ascending order of significance passing the cut-off (adj. p<0.05). (F) The GO analysis of Biological Process and Cellular Component from filtered Miro1-OMM proteomes in wild-type iNs against SNCA-A53T, generated from g:profiler, with the same criteria as in (B, D). (G, H) The volcano plot of prey clusters labeled from Miro1-WT against those from Miro1-OMM in HEK (G) and GBM cells (H). Red- or light blue-colored dots are filtered by p value<0.05 and log_2_FC≥1.0 and grey-colored dots are not significant. (G) Deep blue star-shaped dots are annotated with MitoCarta3.0. Diamond-shaped, orange-colored dots are representative hits from Miro1-WT in HEK293T. (H) Star-shape, light blue- or red-colored dots are representative hits from Miro1-OMM or Miro1-WT in GBM cells, respectively. x axis is log_2_FC between Miro1-WT and Miro1-OMM (right=enriched in Miro1-WT, left=enriched in Miro1-OMM). y axis indicates -log_10_ adjusted p value from the statistical test (higher= more significant). (I) The subcellular localization information from the filtered preys of Miro1-OMM, Miro1-WT, and Miro1-v4 in each cell line. All information is assigned primarily from MitoCarta3.0, organelle IP, and HPA database, manually. Color code is assigned from Prism 10. (J) The depiction of subcellular organelle distributions of filtered Miro1-centered subcellular proteomes. Alpha fold 3 (AF3)-simulated structure of Miro1 (green-colored and localized at mitochondrial surface) and Miro1-v4 (violet-colored and dually localized at mitochondrial surface and peroxisome) are used. All information is assigned primarily from MitoCarta3.0, organelle IP, and HPA database, manually. Grey-colored proteins are from HEK293T, and brown-colored texts are GBM cells.

Notably, GBM cells exhibited fewer cytosolic proteins and a greater number of mitochondrial surface-enriched proteins compared to HEK293T cells (Figure 4A-D), suggesting cell-type-specific differences in subcellular proteome organization. GO analysis revealed that the cytosolic proteomes labeled by EGFP were functionally similar between HEK293T and GBM cells, enriched in expected pathways such as cytosol and nucleoplasm (Figure 4B, D). In contrast, the mitochondrial surface proteomes captured by Miro1-OMM diverged. In HEK293T cells, top-enriched categories were related to mitochondrial membrane organization. In GBM cells, however, the OMM proteome was enriched in proteins associated with extracellular vesicles, organelles, and exosomes (Figure 4B, D). This cell-type-specific difference suggests that the mitochondrial surface proteome is plastic and may adapt to the specialized needs of the cellular environment. To further support this observation, we analyzed whole-cell proteome and found evidence of differential enrichment in pathways involved in metabolic rewiring, cell-cell contact, and cytoskeletal remodeling in GBM cells (Figure S4C-E, Table S3). These data imply that cancer cells may reconfigure their mitochondrial surface to facilitate cell-cell communication and adapt to altered bioenergetic demands.

We next compared the Miro1-OMM proximity-labeled proteome in human induced neurons (iNs) with or without a PD mutation (*SNCA-A53T*). Strikingly, *SNCA-A53T* neurons exhibited a substantial reduction in the number and abundance of proteins labeled by the Miro1-OMM bait compared to wide-type neurons (Figure 4E-F). This loss of mitochondrial surface-associated proteins may reflect the mitochondrial dysfunction known to occur in PD models, and suggests that A53T-driven mitochondrial defects extend to disruptions in surface interactomes, an area that warrants further investigation. Together, our data show that the mitochondrial surface proteome is not static but exhibits considerable plasticity across cell types.

### Mitochondrial surface baits reveal spatially resolved and dynamic interactions with other compartments

Although Miro1-WT and Miro1-OMM share the same C-terminal TM domain anchoring them into the OMM, they differ in stoichiometry and subcellular exposure. Miro1-WT contains cytosolic GTPase and EF-hand domains, while Miro1-OMM lacks these, potentially alternating its interaction radius. To test whether these structural differences led to spatially distinct proximity labeling patterns, we compared their proteomic profiles across GBM and HEK293T cells. As expected, Miro1-WT and Miro1-OMM captured overlapping but also distinct sets of preys in both cell types (Figure 4G-H). GO analysis revealed shared but also specific functional modules enriched in each bait (Figure S5A-B), indicating that altering bait structure can resolve spatially separated subdomains of the mitochondrial surface proteome. Notably, proteins involved in translation and ribosomes were differentially enriched in Miro1-WT bait (Figure S5A-B; discussed later), suggesting that Miro1’s cytosolic domains define distinct interaction landscapes. Importantly, both Miro1-WT and Miro1-OMM baits enriched for proteins localized to multiple organelles, including the ER, nucleus, plasma membrane, Golgi apparatus, and stress granules (Figure 4I-J; S5C-G), displaying extensive inter-organelle connectivity at the mitochondrial surface. Many of these interactions were not detected in previous studies using similar baits, such as Miro2-based BioID ^4^. Although Miro1 and Miro2 share similar membrane-anchoring domains, the proximity labeling method plays a critical role in shaping the detectable proximity interactome. In particular, LOV-Tb enables precise temporal control with minimal background, allowing the detection of transient, low-affinity, or stimulus-responsive interactions that may be missed by the slower, constitutively active BioID. Conversely, BioID may preferentially label more stable and long-lived interactions such as mitochondria-ER anchoring proteins.

Together, these findings demonstrate that our light-activated PL approach enables high spatiotemporal resolution, revealing specialized and previously underappreciated protein assemblies at the mitochondrial surface and their dynamic contacts with extra-mitochondrial compartments.

### Miro1 proximally interacts with ribosomes and TOM complexes at the OMM

Analysis of our PL datasets consistently revealed the presence of ribosomal subunits and translation-associated proteins near the OMM (Figure 3, 4, 5A, S5A-B). Further sub-organelle localization analysis showed that over 50% of MitoCarta3.0-annotated preys identified in our datasets were localized to internal mitochondrial compartments. Notably, proteins with shorter coding sequences (CDS<200 amino acids) tended to localize to the IMS, while those with longer CDSs (>200 amino acids) were predominantly targeted to the matrix (Figure 5B-C, S5H). Recent observations showed that nuclear-encoded intra-mitochondrial proteins often require co-translational import ^34–36^– nascent polypeptides emerge from ribosomes at the OMM and engage the TOM complex. Since all LOV-Tb-tagged Miro1 baits face the cytosol and are not expected to label resident intra-mitochondrial proteins, we reasoned that our labeling may have occurred during co-translational import, or post-translationally after nascent polypeptides exit the ribosomes heading to the TOM complex for import. The fast kinetics of LOV-Tb support its ability to label such transient intermediates. Indeed, major components of both large and small ribosomal subunits were detected as Miro1-WT proximity interactors (Figure 5D, Table S2). Our dataset also revealed close spatial association between Miro1-WT and multiple TOM complex subunits (Figure 1, 5D, Table S2). Importantly, we did not detect mitochondrially encoded proteins, whose translation occurs entirely within the mitochondria, further supporting the conclusion that the labeled peptides are nuclear-encoded and surface-accessible during import. Those findings suggest a possible function of Miro1 in surface protein biogenesis or import.

**Figure 5.**
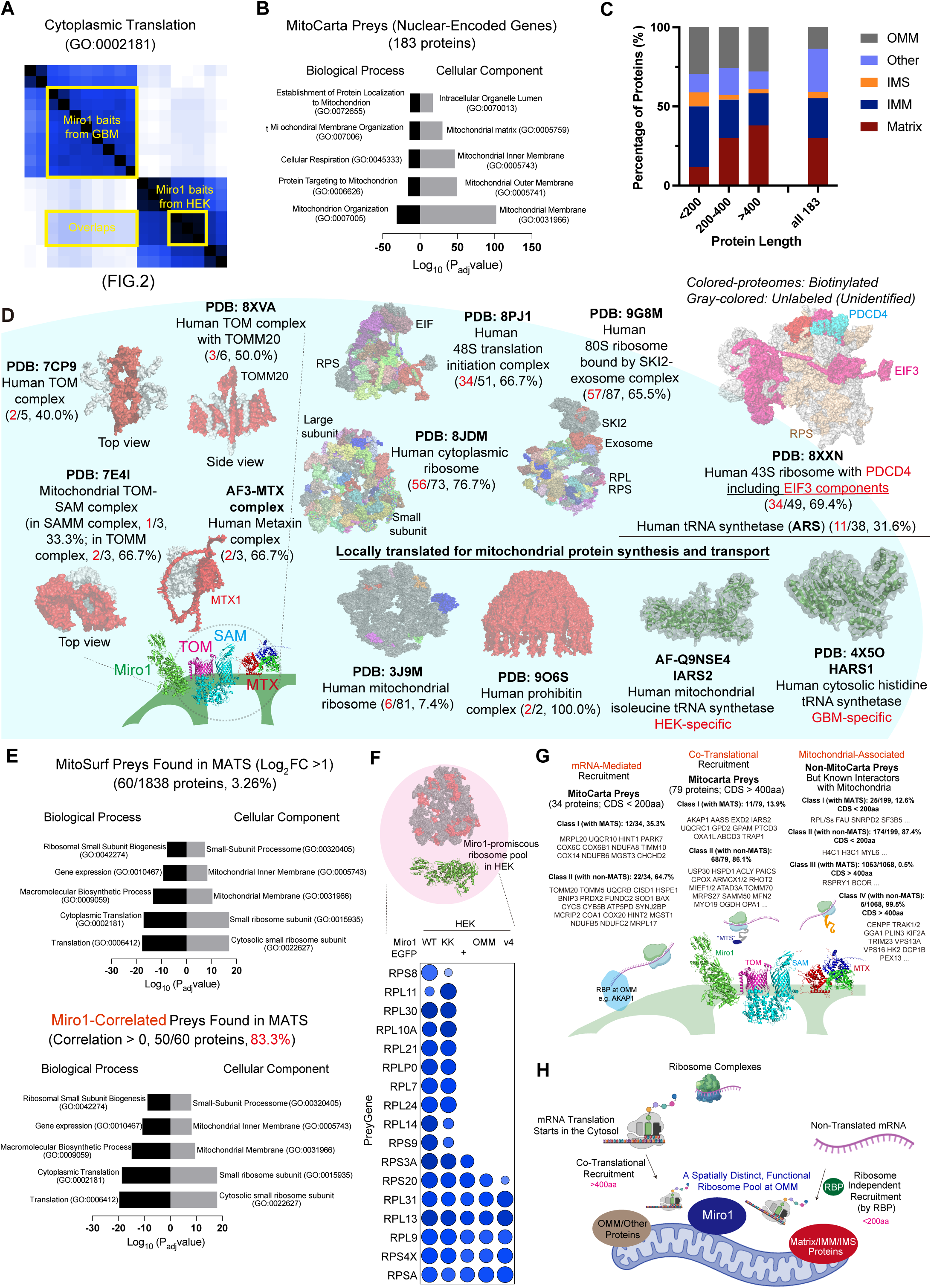
Miro1 proximally interacts with ribosome complexes, TOM, and intra-mitochondrial proteins. (A) A specific GO term (Cytoplasmic Translation, GO:0002181) is highlighted in the same heatmap as in Figure 2A and is enriched in the Miro1-derived baits. (B) The GO analysis from 183 filtered MitoCarta preys. GO terms are significantly enriched in the dataset generated from Enrichr (https://maayanlab.cloud/Enrichr/). (C) Bar graph shows the percentage of proteins from Figure 5B in different sub-mitochondrial compartments, categorized by protein length. (D) The complex-wise distribution of the selected proteomes into structured complexes by Cryo-EM or Alphafold3 (AF3) structures. The biotinylated proteomes from our dataset are colored on the structures from TOM/SAM complex (PDB: 8XVA, 7CP9, 7E4I), AF3-simulated metaxin complex, human cytoplasmic ribosome (PDB: 8JDM), human 48S translation initiation complex (8PJ1), human 80S ribosome bound by SKI2-exosome complex (9G8M), and human mitochondrial ribosome complex (3J9M). Grey-colored means not identified. (E) The GO analysis of 60 filtered MitoSurf preys (upper; Figure S5I) and 50 filtered Miro1-correlated MitoSurf preys (lower), overlapped with MATS, generated from Enrichr. (F) The complex-wise distribution and dot plot profiles of Miro1-proximal ribosome pool in HEK cells. The Cryo-EM structure of cytoplasmic ribosome (PDB: 8JDM) from (D) was used. Red colored means labeled components by Miro1-WT. (G) Classification of Miro1-WT-proximal proteins based on mitochondrial annotation, CDS length, and predicted targeting signals. (H) Our model depicting a Miro1-organized translation compartment at the OMM. RBP, RNA-binding protein.

### RiboLOOM and MitoSurf uncover a spatially distinct, functional ribosome pool at the OMM

The ability of our baits to detect transient, dynamic movements of nascent peptides as they depart ribosomes provides a means to further dissect OMM protein biogenesis. Although recent studies have shown that a subset of mRNAs, including those encoding mitochondrial proteins, are translated by ribosomes localized at the mitochondrial surface ^34–37^, it remains unclear how these ribosomes are recruited, assembled, or stabilized there. When we compared our MitoSurf preys to a mitochondria-associated translation set (MATS) – actively translated mRNAs enriched at the mitochondrial surface detected by Ribo-Seq in a recent study (LOCL-TL) ^34^ (Figure S5I), we discovered that the overlapped hits, potentially proteins being actively translated at the OMM, dominated GO pathways of cytosolic ribosomal subunits and translation among Miro1-WT-correlated preys (Figure 5E). This finding mirrors the earlier observation showing enhanced presence of translation and ribosome proteins in proximity to Miro1-WT bait (Figure S5A-B). When we examined individual ribosome subunits, we discovered cell-type-dependent, selective loss of ribosome components by Miro1-KK, Miro1-OMM, or Miro1-v4 as compared to Miro1-WT (Figure 5F, Table S2). These findings suggest that the cytosolic domain of Miro1 contributes to the organization of OMM-associated ribosome complexes and local translation. Moreover, the putative nascent, mitochondria-targeted polypeptides detected near Miro1-WT included both short and long CDS (Figure 5C, G, S5H, Table S2), corresponding to mitochondrial proteins whose mRNA recruitment mechanisms to the mitochondrial surface are thought to differ ^34^. We therefore propose that ribosomes at the OMM engaging either long CDS or newly recruited short CDS-containing mRNAs represent a distinct functional pool that is maintained by Miro1, forming a spatially confined translation compartment for mitochondrial protein biogenesis (Figure 5H).

To test this hypothesis, we examined preys labeled by Miro1-KK, one of the Miro1-derived baits that lost specific ribosome subunits compared to Miro1-WT (Figure 5F, Table S2). Among the Miro1 variants, Miro1-KK represents the most conservative mutation, preserving the overall structure and domain organization of the wild-type protein while substituting only two residues in the EF-hand motifs that eliminate Ca²⁺ binding and associated conformational changes ^38,39^. Quantitative analysis of proximity-labeled proteins revealed both unique and shared proteins between HEK293T and GBM cells, enriched in Miro1-WT or in Miro1-KK bait (Figure S6A). Because Miro1-KK altered ribosome composition in proximity to Miro1-WT (Figure 5F, Table S2), we reasoned that some of the proteins reduced in Miro1-KK may reflect diminished local translation. Indeed, comparison of proteins decreased in Miro1-KK with MATS, the database of actively translated mRNAs at the OMM ^34^, identified a subset of locally translated preys (Figure 6A-B). Among them was MRPL20, a nuclear-encoded mitochondrial ribosome subunit, that was significantly reduced in Miro1-KK in both HEK293T and GBM cells (Figure 6A-D). These data suggest that loss of ribosome subunits may affect local protein translation at the OMM in Miro1-KK.

**Figure 6.**
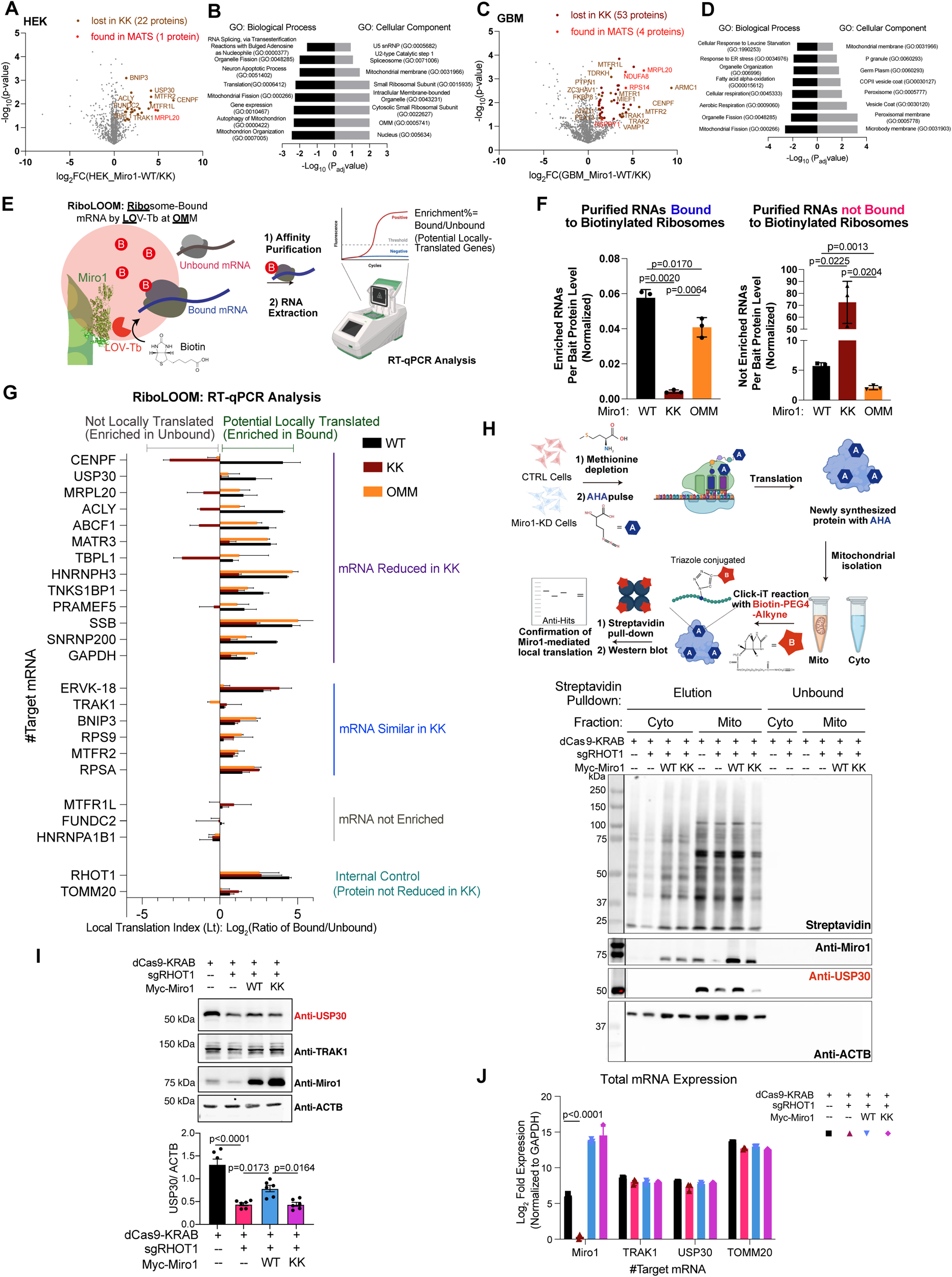
MitoSurf and RiboLOOM uncover a spatially distinct, functional ribosome pool. (A, C) Volcano plots of prey clusters labeled from Miro1-WT against those from KK in HEK293T (A), and GBM (C). Filtration condition is p<0.05 and log_2_FC≥1.0. (B, D) The annotated GO terms from filtered preys in (A, C), generated from Enrichr. Bar length is proportional to the enrichment statistics (-log_10_ adjusted p value); longer bars indicate more strongly enriched terms. The bars are sorted in ascending order of significance passing the cut-off (adj. p<0.05). (E) The schematic illustration of Miro1-centered RiboLOOM. (F) Quantification of purified RNAs associated with biotinylated polysomes (left) versus those not bound (right) in cells stably expressing Miro1-WT, Miro1-KK, or Miro1-OMM. Amounts of RNAs were normalized to bait expression levels. n=3. (G) RiboLOOM targeted RT-qPCR analysis. For each transcript, the local translation index (Lt) is plotted as log_2_(ratio of RNAs from bound versus unbound fraction) in cells stably expressing Miro1-WT (Black), -KK (Red), or -OMM (Orange). Positive value (avg>0.2) (right-side) indicates mRNAs enriched in the bound fraction, whereas negative values (left-side) indicate transcripts preferentially in the unbound fraction. n=3. (H) AHA assay. Newly synthesized proteins were immunoblotted as indicated. (I) Representative immunoblots as indicated and quantification of USP30 bands normalized to ACTB. n=6. (J) Total mRNA expression of the indicated targets measured by RT-qPCR, normalized to GAPDH and expressed as log_2_FC. n=3. One-Way Anova PostHoc Tukey Test (F, I, J).

Because the MATS dataset was generated using a different OMM bait (OMP25) and labeling enzyme (BirA) ^34^, which may define a spatially distinct set of OMM-associated mRNAs, we hypothesized that additional preys lost in Miro1-KK but absent from MATS could represent transcripts translated specifically near Miro1. To identify such mRNAs bound to the Miro1-organized ribosome pool, we established a platform termed RiboLOOM (**<u>Rib</u>**osome-bound mRNAs by **<u>LO</u>**V-Turbo at the **<u>OM</u>**M), which maps Miro1-dependent ribosome-associated mRNAs at the mitochondrial surface (Figure 6E). Quantitative analysis of total mRNAs normalized to bait abundance revealed significant reductions in ribosome-bound transcripts labeled by Miro1-KK or Miro1-OMM compared with Miro1-WT (Figure 6F, S6B), consistent with the decreased proximity of selective ribosomal subunits observed for these baits (Figure 5F, Table S2). However, the loss of ribosome-bound mRNAs was not due to global depletion of ribosomal proteins or the RNA-binding factor AKAP1 ^34^ (Figure S6C-D, Table S2), suggesting that defective ribosome assembly or tethering underlies this reduction.

Targeted reverse transcription quantitative polymerase chain reaction (RT-qPCR) of mRNAs encoding the 23 proteins diminished in Miro1-KK HEK293T cells (Figure 6A, Table S4) confirmed robust enrichment of at least 19 in Miro1-WT-labeled ribosomal fractions relative to unbound controls (Figure 6G), indicating that these proteins may be locally translated near Miro1. This enrichment of both long-CDS (e.g., CENPF, USP30, ACLY) and short-CDS (e.g., MRPL20) mRNAs was markedly reduced in Miro1-KK or Miro1-OMM cells (Figure 6G). Importantly, MRPL20 mRNAs were previously detected in MATS and MRPL20 protein was lost in Miro1-KK PL data (Figure 6A-D), validating the reliability of our assay. In contrast, mRNAs of TRAK1, ERVK-18, BNIP3, MTFR2, RPS9, or RPSA were unchanged despite loss of their proteins in Miro1-KK, suggesting that the protein decrease arises from interaction failure rather than reduced translation. Together, these data identify a distinct pool of mRNAs bound to ribosomes in proximity to Miro1.

To prove mRNAs bound to Miro1-organized ribosomes were being translated locally, we performed a mitochondria-targeted L-Azidohomoalanine (AHA) assay, which detects newly translated proteins by incorporating alkyne-biotin tagged AHA into actively synthesized peptides via Click-iT (Figure 6H). Depletion of Miro1, using dCas9-CRISPRi with non-CDS-targeting gRNAs (Miro1 KD; Figure S6E-I), noticeably reduced mitochondria-localized translation of USP30 (Figure 6H), a top candidate identified in Miro1 RiboLOOM (Figure 6A-G). Miro1-WT, but not Miro1-KK, restored its local translation (Figure 6H). These data mirror the differential enrichment of USP30 mRNAs (Figure 6G) and proteins (Figure 6A-D, Table S4) in Miro1-WT versus KK-labeled ribosomal and protein fractions, consistent with USP30 being locally translated in the Miro1-organized ribosome pool. Removing Miro1 also significantly reduced total protein levels of USP30, which could be partially reestablished by Miro1-WT, but not Miro1-KK (Figure 6I). Crucially, although total protein levels were lower, the corresponding total mRNA levels of USP30 were not decreased in Miro1 KD cells and remained unchanged upon expression of Miro1-KK (Figure 6J). Considering the previously observed reduction in ribosome-bound USP30 mRNAs labeled by Miro1-KK in RiboLOOM (Figure 6F-G), these findings suggest that OMM-associated ribosomes and mRNAs are required for maintaining specific mitochondria-associated proteins, thereby defining their global proteostasis as a spatially regulated process.

Together, these data support our model (Figure 5H) in which disruption of Miro1-mediated ribosome organization at the mitochondrial surface diminishes the engagement of mRNAs and consequently reduces their translation.

### Miro1 is essential for mitochondrial functional integrity

As expected, Miro1 KD HEK293T cells displayed reduced mitochondrial membrane potential detected by TMRM normalized to MitoTracker Green (Figure S6J), altered mitophagy rates (basal and depolarization-triggered) by a ratiometric mitophagy reporter, Mito-mKeima ^40^ (Figure S6K-L), impaired mitochondrial oxygen consumption (OCR), and extracellular acidification rate (ECAR) by Seahorse assay (Figure S6M-N), consistent with the lack of mitochondria-related proteins, e.g. mitochondrial ribosomes and mitophagy players. Miro1-WT which restored OMM protein biogenesis (Figure 6), but not Miro1-KK, significantly rescued some of these defects (Figure S6J-N), supporting the conclusion that Miro1-organized, surface-confined ribosomes are functionally linked to the output of the mitochondria-associated proteome. This newly identified role of Miro1 provides a plausible explanation for the mitochondrial phenotypes, including defects in bioenergetics and crista organization, observed across multiple Miro1 loss-of-function models ^41–44^ (Figure S6J-N), which cannot be fully accounted for by impaired mitochondrial motility alone.

### Pharmacological modulation of the Miro1 proximity proteome

To explore pharmacological regulation of the mitochondrial surface proteome, we applied MR3, a small molecule that binds to the C-terminal GTPase domain of Miro1 (Figure 7A) ^20,28,29,45^. We and others have shown that MR3 confers protective effects in models of disease and aging, without inducing toxicity in healthy cells ^20,28,29,45,46^. The mechanism by which MR3 exerts these effects remains incompletely understood, but we hypothesized that Miro1’s proximity interactions might be involved. To test this, we treated GBM and HEK293T cells expressing Miro1-WT bait with MR3 and performed light-activated PL. We identified subsets of proteins whose enrichment increased or decreased following MR3 treatment (Figure 7B-D, S7). While some altered interactors were shared between cell types, many were cell-type-specific (Figure 7D, S7), highlighting a context-dependent Miro1 interactome under pharmacological intervention. Interestingly, MR3 treatment in both GBM and HEK293T cells led to the association of new proteins with Miro1, which participate in various cellular functions (Figure 7E). In our prior studies, MR3 was shown to promote proteasomal degradation of Miro1 following mitochondrial depolarization ^20,28,29^. Consistent with this, we observed increased proximity of ubiquitin-proteasome pathway components (Figure 7E-F), suggesting that MR3 may facilitate recruitment of Miro1 to the proteasome.

**Figure 7.**
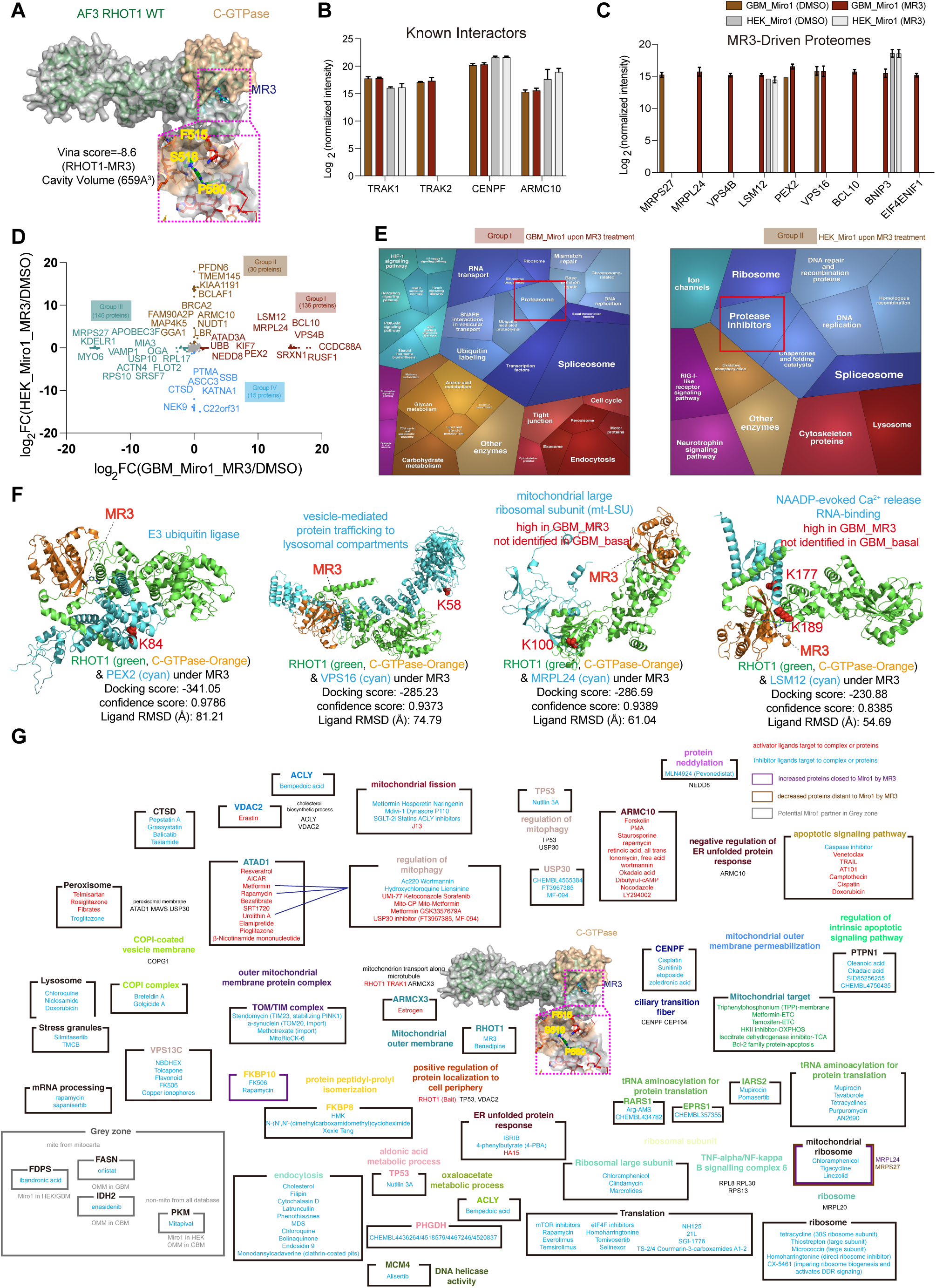
Pharmacological modulation of Miro1 proximity interactome. (A) The structural docking prediction of AF3 Miro1 bounded with the ligand (MR3) and its possible binding residues. (B-C) The MS intensity of known-interacting proteins (TRAK1/2, ARMC10, and CENPF) with Miro1 (B) and of additional proteins (C) by MR3 across cell types. (D) The dual-comparison proteome map by a correlation scatter plot. Each point is a protein with the log_2_FC measured for Miro1-WT, DMSO versus MR3, in two cell models. x axis is log_2_FC (GBM: Miro1-WT, MR3/DMSO) and y-axis is log_2_FC (HEK: Miro1-WT, MR3/DMSO). Positive values indicate higher levels in MR3-treated samples and negatives indicate enrichment in untreated samples (DMSO). Proteins are partitioned into four clustered (Group I-IV) based on their paired fold change pattern: >1 or <-1. Colors of the protein labels match the group colors and grey-colored proteins are not selected: Group I (brick-red, 136 proteins, enriched in GBM, MR3 treated), Group II (brown, 30 proteins, in HEK293T, MR3 treated), Group III (muted-teal, 146 proteins, in GBM, untreated), Group IV (light-blue, 15 proteins, in HEK293T, untreated). (E) Functional category analysis by Proteomaps for new proteins upon MR3 treatment. The size of each polygon correlates with the log_2_FC (+/− MR3). (F) Drug-Miro1 network. Miro1-associated proteins are from Figure 3E. The activator ligands (red), inhibitor ligands (blue), or other ligands (grey) targeting those proteins are shown.

We next examined whether MR3 broadly affected protein abundance beyond Miro1 interactions. Using DDA mass spectrometry, we profiled whole-cell proteome in a panel of cell types: GBM, HEK293T, iPSCs (*SNCA-A53T* and its isogenic control), and iPSC-induced dopaminergic neurons (iDAs) (Figure 4A-B) (Sainz et al., unpublished data). Baseline proteomic differences between these lines confirmed distinct cellular states and lineage-specific programs (Figure S4B), which support our earlier observations of cell-type plasticity of the mitochondrial surface proteome. Notably, MR3 treatment did not induce global proteomic changes in any of the cell types tested or protein level changes of preys whose distances to Miro1 were altered by MR3 (Figure S4B, F). Thus, our data demonstrate that MR3 selectively modulates Miro1-protein proximity interactions in a cell-type-dependent manner, without broadly perturbing proteome composition.

### Drug-protein network identifies mitochondrial surface protein targets

We wondered whether Miro1 proximity interactors from our dataset could be similarly targeted by small molecules. To this end, we profiled known FDA-approved drugs that could bind to Miro1-associated proteins (Figure 4) in our database. Like MR3, those drugs may interfere with Miro1 interaction with those proteins of interest, and our query may identify disease targets on the mitochondrial surface and their therapeutics (Figure 7G). Of all the human targets (80 in GBM, 116 in HEK) that were potentially proximity interactors of Miro1 with a high-confidence score (Figure 4G, H), 50 had 162 drugs or molecules that modulated them and could be overlaid on our protein-interaction network (Figure 7G). Together, we provide a comprehensive protein-drug network for future disease targeting at the mitochondrial surface.

## Discussion

Mitochondria are increasingly recognized as signaling platforms that integrate bioenergetic status, stress responses, and inter-organelle communication. Our study establishes a light-activated PL platform, LOV-Tb-Miro1, to map mitochondrial surface proteomes with high spatial and temporal resolution. Using this tool, we define **MitoSurf**, a resource that reveals spatially resolved and cell-type-specific mitochondrial surface proteomes. We further develop **RiboLOOM**, combining LOV-Tb and ribosome-mRNA analysis, and uncover a spatially distinct ribosome reservoir at the OMM organized by Miro1. This localized ribosome population engages mitochondria-associated mRNAs and is essential for maintaining mitochondrial proteome integrity.

Our findings expand the role of Miro1 beyond its well-established functions in mitochondrial motility and quality control ^20,22,28,29,39,45,47–51^. Previous studies have characterized mechanisms for mRNA recruitment to the mitochondrial surface and defined subsets of transcripts translated near the OMM ^34^. However, how ribosomes are organized and maintained at this site, and how their location contributes to localized and whole-cell protein synthesis, remained unknown. We show that Miro1 proximally interacts with ribosomes and TOM complex components at the OMM, positioning ribosomes to interface directly with the import machinery. The loss of ribosomal subunits or actively translated mRNAs in Miro1-KK, Miro1-OMM, or Miro1-v4 baits demonstrates that Miro1 scaffolds a stable, functional ribosome pool necessary for localized translation. This finding identifies a downstream step following mRNA recruitment, the organization and stabilization of OMM-associated ribosomes, that enables spatially confined translation of nuclear-encoded mitochondrial and other proteins.

Functionally, our data suggest that ribosomes organized by Miro1 are not fully interchangeable with the bulk cytosolic pool. When OMM-associated ribosome components are displaced, total protein output of USP30 declines even though total mRNA levels remain unchanged (Figure 6), indicating that translation of a subset of proteins is spatially restricted to the OMM. This spatial dependency offers a mechanistic explanation for diverse phenotypes observed in Miro1 loss-of-function models ^41–44^ (Figure S6J-N). Our study proposes that Miro1 integrates motility, calcium signaling, quality control, and localized translation to coordinate mitochondrial homeostasis dynamically.

This newly identified function of Miro1 complements previously characterized mitochondrial processes. For example, we and others have shown that Miro1 is targeted by PINK1 and Parkin for proteasomal degradation prior to mitophagy following mitochondrial depolarization ^22^. The removal of Miro1 not only halts mitochondrial motility, thereby segregating damaged mitochondria from the healthy network ^22^, but, as we now demonstrate, also promotes the disassembly of mitochondria-associated ribosomes to prevent continued translation at the site of damage. In aging and disease contexts characterized by resistance of Miro1 to removal, the resulting impairment of mitophagy ^20,28,29,45,49–51^, together with dysregulated local and global translation, may act in concert to promote cellular pathology.

Our approach also provides a blueprint for dissecting dynamic RNA-protein-organelle interfaces. The light-gated nature of LOV-Tb enables temporal control, allowing us to capture transient events such as ribosome engagement, nascent chain elongation, and import coupling. RiboLOOM extends this capability to define ribosome-associated mRNAs in space and time, offering a versatile tool for studying translation at other organelle surfaces. Combining these technologies with genetic and pharmacological manipulations, will open the door to understanding how mitochondrial surface translation adapts to stress, signaling, and disease.

Beyond mitochondria, local ribosome anchoring could represent a general mechanism for maintaining organelle identity and function through compartmentalized translation. Future work integrating LOV-Tb-based proteomics, RiboLOOM-transcriptomics, and super resolution microscopy may illuminate the ultrastructural organization of these localized translation systems. This foundational principle of organelle-anchored translation platforms may govern subcellular autonomy and global translation, which may deteriorate in aging and disease.

## Author Contributions

C.S.K. designed and performed experiments, analyzed data, and made figures. Z.D. built the website. J.S.C., E.M.W., M.B.M., J.E.E. designed and conducted mass spec experiments. X.W. conceived and supervised the project with input from all authors. C.S.K. and X.W. wrote the paper with assistance from all authors.

## Acknowledgements

We thank Drs. A. Ting, M. Lim, A. Venida, and M. Bassik for providing reagents, A. S. Durairaj for technical support, Drs. A. Ting and M. Bassik for discussion, the Vincent Coates Foundation Mass Spectrometry Laboratory, Stanford University Mass Spectrometry (SUMS; RRID:SCR_017801) for processing mass spec samples, and the following funders: National Institutes of Health (RO1NS128040 and RO1GM143258, X.W.; P30CA124435 and 1S10OD030473, SUMS), Stanford Bio-X Seed Grant (X.W.).

## Methods

### EXPERIMENTAL MODEL AND STUDY PARTICIPANT DETAILS

#### Human cells and tissues

No human subjects were used in this study. The iPSC work was approved by Stanford Stem Cell Oversight Committee. HEK293T/17 and GBM (T98G) cells were obtained from ATCC. Details of cell culture conditions and authentication are in method details.

### METHOD DETAILS

#### Constructs, sub-cloning, and plasmid DNA generation

pRK5-Myc-Miro1 (WT and KK) were described in ^39^. Plasmid DNA constructs were cloned using in-house 2x Gibson assembly master mix (2 U T5 exonuclease (M0363S, NEB), 12.5 U Phusion polymerase (M0530S, NEB), and 200 U Taq ligase (M0208L, NEB), ISO buffer containing 1 M Tris pH 7.5, 2 M Magnesium chloride, dNTP (N0447S, NEB), PEG-8000 (V3011, Promega), and 100 mM NAD+ (B9007S, NEB) with PCR-amplified insert genes (or annealed double stranded DNA oligos) and restriction enzyme-digested backbone vectors. The Gibson-assembled DNA plasmids were transformed into competent bacterial cells (XL-1 Blue, 200249, Agilent; gift from Alice Y. Ting at Stanford University) and sequenced after mini-prep (silica membrane mini spin column, 1910-250, Epoch). For high efficiency transformation of the plasmid DNA products from the Gibson assembly, the competent bacterial cells were generated using Mix & Go competent cells transformation kit (T3001, Zymo) according to their instructions. The competent cells are maintained for high efficiency of transformation (about 99% efficiency for 1pg of plasmid DNA). All plasmid DNA constructs were confirmed from whole plasmid DNA sequencing by ELIM Biopharmaceuticals, Inc. (Hayward, CA). Detailed information of all constructs in this study is Table S1.

#### Cell culture

HEK293T/17 (CRL-11268, ATCC) or T98G (CRL-1690, ATCC) cells were maintained in Dulbecco’s modified Eagle’s medium (DMEM, GIBCO) or ATCC-formulated Eagle’s Minimum medium (EMEM, 30-2003, ATCC) supplemented with 10% (v/v) fetal bovine serum (FBS) (35-011-CV, Corning, heat-inactivated) and 1% penicillin/streptomycin solution (15140122, Invitrogen) at 37 °C in a 5% CO_2_ humidified incubator. iPSCs engineered with *NGN2* from a healthy 30-year-old, male donor (https://www.coriell.org/0/Sections/Search/Sample_Detail.aspx?Ref=GM25256) (WT) and edited with *SNCA-A53T* in the WT background (Sainz et al., unpublished) were maintained in mTeSR1 plus (100-0276, STEMCELL) supplemented with its corresponding supplement solution at 37°C in a 5% CO_2_ humidified incubator. For iPSC passaging, cells were harvested using Accutase (07922, STEMCELL) and replated in mTeSR1 plus containing 10 μl ROCK inhibitor (Y-27632, 72307, STEMCELL).

#### Lentiviral production and stable cell line generation

Lentiviral packaging vectors–psPAX: pMD2.G:pRSV-rev (#12260, #12259, #12253, Addgene) and 1 μg of a lentiviral reporter plasmid (1000: 375: 375: 250 ng) were mixed with 9 μg of PEI (1 mg/ml, Polyscience) in 200 ml DMEM without FBS, and the mixture was added onto HEK293T cells cultured in one well of a 6-well plate in 2 ml of DMEM without FBS at 37°C with 5% CO_2_. After incubation for 24 hours, DMEM containing the DNA-PEI mixture was removed and replaced with 2 ml of prewarmed DMEM, supplemental with 10% FBS. Lentiviral particles were first harvested from the removed 2-ml DMEM. On the next day, lentiviral particles were harvested from the 2-ml of complete DMEM for a second time. The medium containing lentiviruses was filtered with 0.42 mm pore size syringe filter, precipitated using the PEG-8000 Virus Precipitation Solution for at least 4 hours at 4°C, and then centrifuged at 1600 x g for 1 hour at 4°C. After centrifugation, supernatants were gently discarded without disturbing co-precipitated lentiviral particles with PEG-8000, and pellets containing concentrated lentiviral particles were resolubilized in 1ml of fresh PBS (4x concentrated lentiviruses from total 4 ml crude cultured DMEM) and aliquoted (200 μl in flash freeze tubes) to avoid repeated thaw-freeze cycles before being stored in a -80°C freezer. To transduce concentrated lentiviral particle into cells (HEK293T, T98G, and iPSCs), 200 ml of concentrated viral suspension in 1 ml of growth medium was added dropwise to target cells at ≥75% confluent cell density cultured in growth media, incubated for 24 hours before fresh media change at 37°C. At least 48 hours of transduction allowed lentiviruses to integrate into the host genome. Transduced cells were grown for 1 week to enlarge cell population. During this time, cells were validated with immunostaining or Western blotting.

#### CRISPRi construct and *Miro1*-knockdown cell line generation

The plasmid DNA constructs including target sgRHOT1 were cloned into the pLV lentiviral vector containing dCas9-KRAB-T2A-EGFP under UbC promoter and target sgRNA under U6 promoter (71237, Addgene) using standard Gibson assembly (E2621S, New England Biolabs). The complimentary primer set for the sgRNA oligos (Miro1: aggcggagctggcgctgtcc), designed by the online design tool (https://crispr.dbcls.jp/gRNAcalc/?seq=aggcggagctggcgctgtcccgg), with the overhang region (Tm= 55°C), was subcloned into the target vector with BsmBI. The top and bottom strands of oligos were resuspend to a final concentration of 100 μM and annealed in a thermocycler by using the following parameters: 37°C for 30 minutes; 95°C for 5 minutes; ramp down to 25°C at 5°C per minute. Next, annealed oligos were dilute to 1:200 with clean water. The digested and cleaned vector with BsmBI and the annealed sgRNA oligos were assembled by standard Gibson assembly master mix at 55°C for 45 minutes in a thermocycler and transformed into competent bacterial cells (DH5alpha, 18258012, Invitrogen) and sequenced after mini-prep (1910-250, Epoch). To generate a dCas9-KRAB stable cell line, cells were lentivirally transduced with a vector encoding dCas9-KRAB fusion protein conjugated with T2a-EGFP as a fluorescent marker under UbC promoter and sgRHOT1 under U6 promoter. Two days after the transduction, single cell with EGFP is selected from a 96-well culture plate. The efficiency of the CRISPRi system in repressing the target gene, *Miro1*, was validated using RT-qPCR and Western blotting. Forward Primer: ttttaacttgctatttctagctctaaaacGGACAGCGCCAGCTCCGCCTggtgtttcgtcctttccac; Reverse Primer: gtggaaaggacgaaacaccAGGCGGAGCTGGCGCTGTCCgttttagagctagaaatagcaagttaaaa

#### iN differentiation

Human iPSCs were transduced as described above. iPSCs were differentiated into cortical neurons according to a previously established protocol ^52^. Briefly, iPSCs were dissociated using accutase, and 700,000 cells were plated onto a Matrigel-coated 6-well plate in pre-differentiation medium. This medium consisted of DMEM/F12 (11330032, Gibco) with 2 μg/ml doxycycline (D3447, Sigma), 1% N2-supplement (17502048, Gibco), 1% NEAA (11140050, Gibco), 1% GlutaMAX (35050061, Gibco), and 10 μM ROCK inhibitor (1254, Tocris). The media was refreshed after 24 and 48 hours with and without rock inhibitor respectively. At 72 hours, the pre-differentiated iNs were dissociated with accutase and 40,000 cells were seeded onto poly-L-ornithine (P4957, Sigma) coated coverslips (PCS-1.5-10, Mattek) in a 24-well plate. The neurons were then cultured in BrainPhys Neuronal Medium (05790, STEMCELL), supplemented with 2% B27 (17504-044, Gibco), 10 ng/ml BDNF (450-02, PeproTech), 10 ng/ml NT-3 (450-03, PeproTech), and 1 μg/ml Laminin (23017-015, Thermo). On day 5 and 7, iNs were utilized for the experiments.

#### iDA differentiation

Human iPSCs were differentiated into iDAs using a previously established protocol ^53^. Briefly, iPSCs were plated on Matrigel-coated 6-well plates at a density of 6x10^5^ cells/well in mTesr1 medium supplemented with Rock Inhibitor (10 μM). The next day, media was replaced with DMEM/F12 (10565-018, Thermo) supplemented with 1x N2 (17502-048, Thermo), 1x B27 (17504-044, Thermo), 1x non-essential amino acids (11140-050, Thermo), 10 ng/mL BDNF (450-02, Peprotech), 10 ng/mL GDNF (450-10, Preprotech) and 2 μg/ml doxycycline (D3447, Sigma) (Day 0/1 media). Forty-eight hours later, cells were dissociated with accustase and seeded on plates or coverslips coated with PDL (30 minutes at room temperature, followed by 3 washes with sterile water and allowed to dry for 2 hours) and Laminin (10 μg/ml added to plates or coverslips following PDL coating; incubated at 37°C for two hours prior to cell seeding) (23017-015, Thermo). Cells were seeded in STEMDiff Midbrain differentiation medium (100-0038, STEMCELL) supplemented with 2 μg/mL of Doxycycline, 1 μg/mL mouse laminin and 200 ng/mL of SHH (78065, STEMCELL). Full media changes were performed from days in vitro (DIV) 3-4, and half media changes were conducted from days 5-8 with supplemented differentiation medium. On DIV 9, media was fully changed to STEMDiff Midbrain Neuron Maturation media (100-0041, STEMCELL) supplemented with 2 μg/ml doxycycline and 1 μg/ml mouse laminin. Starting on DIV 10 and every two days thereafter, half the media was changed with supplemented maturation medium. Neurons were harvested on DIV 20-21 for experiments. Purity of iDAs was determined by immunocytochemical analysis in Sainz et al., 2026.

#### In situ biotinylation with LOV-Tb for immunoblotting and immunostaining

HEK293T/GBM/iN cells transiently or stably expressing LOV-Tb under a suitable promoter were expressed or induced with 100-200 ng/ml doxycycline overnight under TRE3G promoter. After induction, cells were covered with aluminum foil until ready for blue light stimulation. When uncovered, cells were handled in a dark room with red light illumination to avoid undesired activation of LOV-Tb. To stimulate LOV-Tb, biotin was added to cells to a final concentration of 50 μM and incubated at 37°C while being placed directly on top of a blue light light-emitting diode (LED) array (470 nm). Continuous blue light activates LOV-Tb. Control samples were processed in parallel omitting light or biotin. Following light stimulation, cells were once again handled under red light, washed in ice-cold Dulbecco’s PBS (DPBS) three times, and analyzed by Western blotting or immunofluorescence as described below.

#### Confocal microscopy

For immunostaining, cells were fixed in 4% paraformaldehyde (sc-281692, Santa Cruz Technologies) for 15 minutes at room temperature or fixed in cold methanol for 15 minutes at -20°C and then washed twice in PBS. Cells were permeabilized and blocked in TBS with 5% BSA and 0.3% Triton X-100 for 30 minutes, and then immunostained with following antibodies; rabbit anti-Oct4 (ab19857, abcam), mouse anti-V5 (R960-25, Invitrogen) at 4°C overnight or for 1 hour room temperature, followed by Alexa Flour 488-conjugated goat anti-mouse IgG (A-11001, Invitrogen) at 1:3,000 and Alexa Flour 568-conjugated-Streptavidin antibodies (S11226, Invitrogen) at 1:10,000 at room temperature for 30 minutes. After incubation, cells were washed twice in 1x TBST (TBS with 0.05% Tween-20) and mounted on slides with mounting solution containing DAPI (P36931, Thermo). All samples were imaged at room temperature with a 63x/N.A.1.30 oil Plan-Apochromat objective on a Leica SPE laser scanning confocal microscope with identical imaging parameters. For live cell imaging of MitoTracker Green and TMRM, cells grown on 35-mm cell culture dishes were incubated with 100 nM of MitoTracker Green FM (M7514, Invitrogen) and 25 nM TMRM (tetramethylrhodamine methyl ester, T668, Invitrogen) in the complete DMEM medium for 30 minutes. Cells were washed once with pre-warmed dye-free medium. For live cell imaging of cells stably expressing mito-mKeima with MitoTracker Deep Red, cells grown on 35-mm cell culture dishes were incubated with 100 nM of MitoTracker Deer Red FM (M22426, Invitrogen) in DMEM for 30 min. Cells were washed once with pre-warmed dye-free medium. Live cells were then imaged on a Leica SPE laser scanning confocal microscope with a 63x/water objective and appropriate excitation/emission setting (488 nm excitation laser for MitoTracker Green, 561 nm exciting laser for TMRM and Mito-mKeima, and 635 nm exciting laser for MitoTracker Deer Red), as described previously ^28,29,40,49,50^. Fluorescent intensity was measured by Image J (Fiji). The same imaging setting was applied within each experiment. Images were processed with Leica Application Suite (LAS) X Life Science Microscope Software (v.1.4.6, Leica).

#### Mitophagy assay

Cells stably expressing Mito-mKeima were seeded in a 96-well assay plate (black with clear flat bottom) (Corning, 3340) in 100 μl complete culture medium and allowed to attach overnight. Fresh medium containing 10 μM of CCCP or the equivalent volume of the solvent, DMSO, was applied for 24 hours at 37°C. Immediately before the assay, wells were washed once and incubated in pre-warmed buffer (1X PBS, pH 7.4). The plate was read using a TECAN plate reader (M200pro) with excitation wavelength between 401 and 590 nm (step size of 2 nm) and emission wavelength at 620 nm at 37°C. The mitophagy index was calculated for each well as the excitation ratio (F_586_/F_440_). For CCCP-induced mitophagy, the delta mitophagy index was defined as, “mitophagy index of basal”–“mitophagy index of CCCP”.

#### Immunoblotting

All cultured cells were lysed with cell lysis buffer (87787, Pierce) containing protease inhibitor cocktail (78430, Thermo) and then boiled for 5 minutes before being loaded into 8% or 10% SDS-PAGE. Manually made polyacrylamide gels and Tris-glycine-SDS running buffer were used for electrophoresis. Nitrocellulose membrane (1620115, BioRad) was used in wet transfer with transfer buffer for 2 hours. Transferred membranes were stained with Ponceau S followed by destaining with 1xTBST till the color disappeared. The membranes were blocked in TBST containing 5% skimmed milk for 1 hour at room temperature and then immunoblotted with the following primary antibodies in TBST with 5% skimmed milk at 4°C overnight or room temperature for 1 hour: anti-ACTB (β-actin) (13E5, 4970, Cell Signaling) at 1:1,000, mouse anti-V5 (R960-25, Invitrogen) at 1:10,000, rabbit anti-RHOT1 (A22469, Abclonal), rabbit anti-TOMM20 (D8T4N, Cell Signaling) at 1:1,000, mouse IgG2b anti-His6 antibody (MA1-21315, Invitrogen) at 1:1,000, rabbit anti-USP30 (83240-5RR, Proteintech) at 1:1,000, goat anti-TRAK1 (PAB6727, Abnov) at 1:1,000, and rabbit anti-Myc (9B11, Cell Signaling) at 1:1,000. After washes, the following secondary antibodies were used at 1:3,000 for 30 minutes at room temperature: goat anti-mouse IgG2b AF488 (A-21141, Invitrogen), goat anti-rabbit IgG (H+L)-HRP conjugate (1706515, Bio-Rad), IRDye 680RD (926-68070, LICOR-Bio), IRDye 800CW (926-32211, LICOR-Bio), and Streptavidin-conjugated IRDye 800CW (926-32230, LICOR-Bio). For HRP blotting, SuperSignal West Dura Substrate (34075, Thermo) was used. After washes with TBST, the membranes were scanned using Bio-Rad ChemiDoc MP system. Quantification was carried out with ImageJ (Fiji).

#### RNA extraction and RT-qPCR

Total RNA was extracted from cultured cells using TRIzol (GIBCO) according to the manufacturer’s instructions. Concentrations of total RNA were measured using a Nano Drop (Thermo). One μg of total RNA was then subjected to reverse transcription using the Reverse Transcription Mastermix (RK20428, Abclonal). Next, qPCR was performed using the CFX96 instrument (Bio-Rad) and Universal SYBR Green mastermix (RK21205, Abclonal) following the manufacturer’s instructions. qPCR was analyzed by the CFX Maestro Software, and relative expression level was presented as the ratio of the target gene to the internal standard gene, *GAPDH*, or other controls. Each sample was analyzed using three independent biological repeats. All data were processed with Bio-Rad CFX manager (v. 3.1). All primers were listed in Table S1.

#### Seahorse assay

OCR and ECAR were measured using a Seahorse XF HS Mini-Analyzer (Agilent). Cells were seeded in each well of poly-L lysine-coated Seahorse culture plates (103721-100, Agilent) at 4000 cells per well in 100 μl complete growth medium and allowed to attach overnight. On the day of the assay, cells were washed twice with pre-warmed Seahorse assay medium containing 10 mM glucose, 1 mM sodium pyruvate, and 2 mM L-Glutamine for 1 hour in a non-CO_2_ incubator at 37°C (Seahorse XF DMEM assay medium pack, pH7.4 w/o phenol red, 103680-100, Agilent). The sensor cartridge was pre-hydrated overnight in XF Calibrant at 37°C without CO_2_ and then loaded with the following compounds for a mitochondrial stress test: oligomycin (final 1 μM) in port A, FCCP (final 0.5 μM) in port B, and rotenone/antimycin A (final 0.5 μM each) in port C. OCR and ECAR were measured in cycles of mix (3 minutes), wait (2 minutes), and measure (3 minutes). Cells were sequentially treated with oligomycin (oligo), FCCP, and rotenone/antimycin A (Rot/AA), as indicated in Figure S6. Three measurement cycles were recorded at baseline and after each injection. After the assay, cells were lysed in RIPA buffer and protein concentration was determined by BCA assay. OCR (pmol/min) and ECAR (mpH/min) rates were normalized to protein amounts (μg) per 4000 cells after the assay.

#### Sample preparation of LOV-Tb-derived biotinylated proteomes

Cells were grown in 150 mm (Miro1-WT, KK, v4) or 100 mm culture dishes (EGFP and Miro1-OMM). Stably expressed LOV-Tb constructs in HEK293T, GBM, and iPSCs were induced at 60-80 % confluence before the day of in situ biotinylation. Cells were covered with aluminum foil until ready for blue light stimulation. When uncovered, cells were handled in a dark room with red light illumination to avoid undesired activation of LOV-Tb. Fifty μM of biotin was added and incubated at 37°C while being placed directly on top of a blue light LED array for stimulation of biotinylation by LOV-Tb. The cells were washed three times with “cold” PBS and then lysed with 0.750 ml of 2% SDS in 1X TBS (25 mM Tris, 0.15 M NaCl, pH 7.2; 28358, Thermo) with 1X protease inhibitor cocktail and 1 μl of benzonase. For removal of free biotin and detergent, 4 times the sample volume of cold acetone (-20°C, 650501, Sigma) was added to each lysate and kept at -20°C for 2-16 hours. Samples were centrifuged at 13,000 x g for 10 minutes at 4°C. Supernatant was removed gently. Four ml of 90% cold acetone-10% 1X TBS were added to the pellet. Samples were vortexed vigorously and kept at -20 °C for 2-16 hours. On the next day, samples were centrifuged at 13,000 x g for 10 minutes at 4°C. Supernatant was removed gently and was allowed to air dry for 3-5 minutes. Pellets were resolubilized with 0.5 ml of 8 M urea (U5378, Sigma,) in 50 mM ammonium bicarbonate (ABC, A6141, Sigma). Concentration of protein was measured by Pierce 660 nm protein assay reagent (22660, Thermo). Samples were denatured at 650 rpm for 1 hour at 37°C under thermomixer (Eppendorf). The samples were reduced by adding dithiothreitol (DTT) (43816, Sigma) to a 10 mM final concentration and incubated at 650 rpm in a rotator for 1 hour at 37°C. Next, the samples were alkylated by adding iodoacetamide (I1149, Sigma) to a final concentration of 40 mM and incubated under dark condition (covered with aluminum foil) at 650 rpm in a rotator for 1 hour at 37°C. The samples were diluted eight times using 50 mM ABC to dilute urea below < 1 M. CaCl_2_ was added to a 1 mM final concentration to prevent self-cleavage of Trypsin. TPCK-treated Trypsin (20233, Thermo) was added to each sample (50:1 w/w). Samples were incubated at 650 rpm in a rotator for 6-18 hours at 37°C and centrifuged at 17,000 x g for 10 minutes to remove insoluble material. Streptavidin magnetic beads (150 μl or 75 μl for Miro1-derived samples) (88817, Pierce) were washed with 2 M urea in 1x TBS for four times and added to the sample. The samples were rotated for 1 hour at room temperature. The flow-through fraction was kept in -20°C and the beads were washed twice with 2 M urea in 50 mM ABC. To minimize the salts in the samples, the beads were washed with pure water (LC-MS grade) in new 1.5 ml protein lobind tubes. One hundred fifty μl of elution solution (80 % acetonitrile, 900667, Sigma; 0.2 % TFA, T6508, Sigma; 0.1 % formic acid, 28905, Thermo; LC-MS grade water) were added to the beads bound to biotin-modified peptides and heated at 60°C at 650 rpm for 5 minutes. Elution fraction was transferred to new 1.5 ml protein lobind tubes. The elution step from the beads was repeated two more times and all three elution samples were combined. To remove brown-colored beads particles, all samples were centrifuged at full speed for 2 minutes and transferred to new 1.5 ml protein lobind tubes. All elution samples were dried using Speed Vac. The biotin-modified peptide samples were desalted using desalting tips including C18 resin (87782, Thermo). Samples for label-free proteomics were stored at -20°C after the clean-up with the C18 tips until being injected to a mass spectrometer directly. TMT labeling is described below.

#### TMT labeling of biotin-modified peptides

Desalted biotin-modified peptides were labeled with TMT10plex reagents (Thermo). Each TMT tags were solubilized in anhydrous acetonitrile and were allowed to dissolve for 5 minutes with occasional vortexing. Each tube was briefly centrifuged to gather the solution. The biotin-modified peptides were resuspended in 10 μl of acetonitrile and 17 μl of 0.2 M HEPES buffer (pH 8.5) was added to the peptide solution, following labeled with 3 μl of 5 mg/ml TMT10plex reagents. Samples were incubated at room temperature for 1 hour with shaking at 1,000 rpm thermomixer (Eppendorf). TMT reaction was quenched with 2 μl of 5% hydroxylamine solution at room temperature for 15 minutes with shaking. The TMT-labeled biotin-modified samples were combined, dried to completion, reconstituted in 100 μl of 0.1% formic acid, and cleaned up with desalting tips including C18 resin (87782, Thermo).

#### Sample preparation for label-free whole-cell proteomics (iPSCs and iDAs)

Cell pellets of iPSCs and iDAs treated with DMSO and MR3 (10 μM, 24 hours) were lysed on ice with vortexing in 50 μl of 8 M Urea, 75 mM NaCl, 50 mM Tris, pH 8.0, and 1 mM EDTA lysis buffer with protease and phosphatase inhibitors (Thermo). A protein BCA assay (Thermo) was used to quantify protein and 50 μg of each sample was digested. Samples were reduced with 5 mM DTT for 45 minutes at 37°C and then alkylated with 50 mM CAA for 30 minutes at room temperature in the dark. They were then diluted to 1.5 M Urea with 50 mM Tris, pH 8. Samples were then digested overnight for about 15 hours with 1 μg of Trypsin at 800 rpm and 37°C on a ThermoMixer (Thermo). In the next morning, digested samples were quenched with 10% trifluoroacetic acid and desalted on 10 mg Strata-X cartridges (Phenomenex). Eluted peptides were dried. Samples for label-free proteomics were stored at -20°C after the clean-up with the C18 tips until being injected to a mass spectrometer directly.

#### Sample preparation and TMT labeling for whole-cell proteomics (HEK and GBM)

Cell pellets of HEK and GBM were lysed with vortexing in 200 μl of 2% SDS in 1x TBS (25 mM Tris, 0.15 M NaCl, pH 7.2; 28358 Thermo) with 1x protease inhibitor cocktail and 1 μl of benzonase. For removal of detergent in lysates, 4 times the sample volume of cold acetone (-20°C; 650501, Sigma) was added to each sample and kept at -20°C for 2-16 hours. Samples were centrifuged at 13,000 x g for 10 minutes at 4°C. Supernatant was removed gently. Four ml of 90% cold acetone/10% 1xTBS were added to the pellet. Samples were vortexed vigorously and kept at -20°C for 2-16 hours. On the next day, samples were centrifuged at 13,000 x g for 10 minutes at 4°C. Supernatant was removed gently and was allowed to air dry for 3-5 minutes. Pellets were resolubilized with 0.5 ml of 8 M urea (U 5378, Sigma) in 50 mM ABC. Concentration of protein was measured by Pierce 660 nm protein assay reagent (22660, Thermo). Samples were denatured at 650 rpm for 1 hour at 37°C under thermomixer (Eppendorf). The samples were reduced by adding DTT (43816, Sigma) to 10 mM final concentration and incubated at 650 rpm in a rotator for 1 hour at 37°C. Next, the samples were alkylated by adding iodoacetamide (I1149, Sigma) to 40 mM final concentration and incubated under dark condition (covered with aluminum foil) at 650 rpm for 1 hour at 37°C. The samples were diluted eight times using 50 mM ABC to dilute urea below < 1 M. CaCl_2_ was added to 1 mM final concentration to prevent self-cleavage of Trypsin. TPCK-treated Trypsin (20233, Thermo) was added to each sample (50:1 w/w). Samples were incubated at 650 rpm in a rotator for 6-18 hour at 37°C. Samples were centrifuged at 17,000 x g for 10 minutes to remove insoluble material. The digested samples were quenched with 0.1% TFA and desalted with the C18-tips three times. The collected samples were dried using Speed Vac. For TMT-labeling, desalted peptides were labeled with TMT10plex reagents (Thermo). The desalted peptides were quantified with the Quantitative peptide assay (Fluorescence, 23275, Thermo). The peptides (100 ng) were diluted in 10 μl of acetonitrile and 17 μl of 0.2 M HEPES buffer (pH 8.5), followed by being labeled with 3 μl of 5 mg/ml TMT10plex reagents.

Samples were incubated at room temperature for 1 hour with shaking at 1,000 rpm thermomixer (Eppendorf). TMT reaction was quenched with 2 μl of 5% hydroxylamine solution at room temperature for 15 minutes with shaking. The TMT-labeled samples were combined, dried to completion, reconstituted in 100 μl of 0.1% formic acid, and cleaned up with desalting tips including C18 resin (87782, Thermo). The samples were stored at -20°C after the clean-up with the C18 tips until being injected to a mass spectrometer directly.

#### Mass spec data acquisition for whole-cell proteomics (iDAs and iPSCs)

Digested peptides were reconstituted in 45 μl of 98% water, 2% acetonitrile, and 0.1% formic acid, and centrifuged at 15,000 rpm for 15 minutes. Peptide concentration was determined using protein BCA assay working reagents (Thermo) with Peptide Digest Assay Standard (Pierce, Thermo). Each sample was diluted to 0.3 μg/μl and 2 μl was injected by autosampler onto a µPAC™ Trapping Column (Thermo) connected to a 50 cm µPAC™ Neo HPLC Column (Thermo) connected to a PepSep 10 μm liquid junction emitter (Bruker Daltonics, Germany). The column was placed in-line with an Orbitrap Eclipse Tribrid mass spectrometer (Thermo) equipped with a nanoelectrospray ion source (Flex, Thermo) connected in-line to an Ultimate 3000 RSLCnano HPLC system (Thermo). The Orbitrap Eclipse Tribrid mass spectrometer was operated in data dependent mode to automatically switch between MS and MS/MS acquisition with a cycle time of 1 second. Buffer A is 99.9% H_2_O, 0.1% formic acid; buffer B is 99.9% acetonitrile, 0.1% formic acid. The HPLC gradient program delivered an acetonitrile gradient over 60 minutes. For the first 3 minutes, the flow rate was 750 nl/minute at 2% B for the nanopump and 10 ul/minute at 100% A for the loading pump to load the peptides onto the trap column. At 2.8 minutes, the nanopump was changed to 10% B. At 3 minutes, the column switching valve was switched so that the trap column was in-line with the analytical column. At 5 minutes the flow rate was reduced to 300 nl/minute and the solvent composition was changed to 12% B. A two-step gradient was used from 12% to 20% B for 41.8 minutes followed by 20% to 40% B for 15.9 minutes. The flow rate was then increased to 750 nl/minutes for column washing using seesaw gradients up to 90% B and then followed by re-equilibration at 2% B. The MS1 scans were acquired in the Orbitrap at 240K resolution with a 1 x 10^6^ automated gain control (AGC) target, auto max injection time, and a 375-2000 m/z scan range. MS2 targets were filtered for charge states 2-7, with a dynamic exclusion of 60 seconds, and were accumulated using a 0.7 m/z quadrupole isolation window. MS2 scans were performed in the ion trap at a turbo scan rate following higher energy collision dissociation with a 35% normalized collision energy. MS2 scans used a 1 x 10^4^ AGC target and 35 ms max injection time.

#### Mass spec data acquisition for PL-driven proteomics (iNs)

Digested peptides were reconstituted in 14 μl of 98% water, 2% acetonitrile, and 0.1% formic acid and centrifuged at 15,000 rpm for 15 minutes. For each sample, 10 μl was injected by autosampler onto a µPAC™ Trapping Column (Thermo) connected to a 50 cm µPAC™ Neo HPLC Column (Thermo) connected to a PepSep 10 μlm liquid junction emitter (Bruker). The column was placed in-line with an Orbitrap Eclipse Tribrid mass spectrometer (Thermo) equipped with a nanoelectrospray ion source (Flex, Thermo) connected in-line to an Ultimate 3000 RSLCnano HPLC system (Thermo). The Orbitrap Eclipse Tribrid mass spectrometer was operated in data independent mode. Buffer A is 99.9% H_2_O, 0.1% formic acid; buffer B is 99.9% acetonitrile, 0.1% formic acid. The HPLC gradient program delivered an acetonitrile gradient over 120 minutes. For the first 3 minutes, the flow rate was 750 nl/minutes at 2% B for the nanopump and 10 μl/minutes at 100% A for the loading pump to load the peptides onto the trap column. At 2.8 minutes, the nanopump was changed to 10% B and the flow rate reduced to 300 nl/minutes. At 3 minutes, the column switching valve was switched so that the trap column was in-line with the analytical column. A two-step gradient was used from 10% to 20% B for 74 minutes followed by 20% to 40% B for 46 minutes. The flow rate was then increased to 750 nl/minutes for column washing using seesaw gradients up to 90% B and then followed by re-equilibration at 2% B. The MS1 scans were acquired in the Orbitrap at 30K resolution with aa 4 x 10^5^ AGC target, 54 ms max injection time, and a 390-1010 m/z scan range. MS2 scans were performed in the Orbitrap at 30K resolution and 10 m/z window with precursor mass range 400-1000 m/z. Ions were fragmented with higher energy collision dissociation with a 30% normalized collision energy. MS2 scans used a 5 x 10^4^ AGC target, 54 ms max injection time, and 145-1450 scan range (m/z). The scan cycle was set at 3 seconds.

#### Mass spec data acquisition for TMT-labeled peptides for whole-cell and PL-driven proteomics

Proteolytically digested peptides were separated using an in-house pulled and packed reversed phase analytical column (∼25 cm in length, 100 microns of I.D.), with Dr. Maisch 1.9-micron C18 beads as the stationary phase. Separation was performed with a180-minute reverse-phase gradient (2-45% B, followed by a high-B wash) on an Acquity M-Class UPLC system (Waters Corporation, Milford, MA) at a flow rate of 300 nl/minute. Mobile Phase A was 0.2% formic acid in water, while Mobile Phase B was 0.2% formic acid in acetonitrile. Ions were formed by electrospray ionization and analyzed by an Orbitrap Eclipse Tribrid mass spectrometer (Thermo). The mass spectrometer was operated in a data-dependent mode using CID and HCD fragmentation for MS2 and MS3 spectra generation, respectively.

#### Mass spec data acquisition for PL-driven proteomics

Desalted peptides were flash frozen, then dried by centrifugal evaporation, and resuspended in 2% acetonitrile/0.1% TFA. Peptides were analyzed on an Obitrap Fusion Lumos mass spectrometer (Thermo) or a TimsTOF HT mass spectrometer (Bruker) equipped with an EASY-nLC 1200 LC system (Thermo) and nanoFlex ESI source or a nanoElute 2 (Bruker) with an average of 100 ng sample input. For Orbitrap, peptides were separated by capillary reverse phase chromatography on a 25 cm column (75 μm inner diameter, packed with 1.6 μm C18 resin, AUR2-25075C18A, Ionopticks). Electrospray Ionization voltage was set to 2000 volts. Both were operated with a flow rate of 300 nl/minute. Column temperature was maintained at 50°C throughout the procedure. Data was acquired in top speed, Data-Dependent mode with a duty cycle time of 1 second. Full MS scans were acquired in the Orbitrap mass analyzer (FTMS) with a resolution of 120,000 (FWHM) for PL-proteomics. The m/z scan range was 375-1500 m/z. Selected precursor ions were subjected to fragmentation using higher-energy collisional dissociation (HCD) with quadrupole isolation window of 0.7 m/z and normalized collision energy of 31%. HCD fragments were analyzed in the Ion Trap mass analyzer (ITMS) set to Turbo scan rate. Fragmented ions were dynamically excluded from further selection. The maximum injection time was set to Auto for full FTMS scans. For TimsTOF, the injection volume was 8 μl from total 20 μl of resuspension with Evotips. One injection each was shot in Data-Dependent and Data-Independent mode.

#### Mass spec data analysis for pilot samples from PL-derived peptides with D+4 stable isotope-coded biotin

The .d files acquired in data-independent mode were analyzed using Spectronaut v 19.1 (Biognosys AG, Schlieren, Switzerland) with the directDIA+ feature against the Uniprot Homo sapiens protein database. Proteolysis with Trypsin was assumed to be specific, allowing for N-ragged cleavage with up to two missed cleavage sites, with a PSM false discovery rate (FDR) of 1%. Data filtering was based on the Q Value, and global normalization was utilized. Cysteine modified with iodoacetamide was set as a fixed modification. Variable modifications included biotinylation of lysine (mass shift +230.10 Da), oxidation of methionine, deamidation of asparagine and glutamine, deoxidation of methionine and tryptophan, and phosphorylation of serine, threonine, and tyrosine, acetylation of N-terminal and lysine, and pyro-glutamic acid conversion from glutamine and glutamic acid.

#### Mass spec data analysis for label-free quantification of biotinylated peptides (Orbitrap)

MS/MS data were searched by MaxQuant (version 2.7.5.0) with Andromeda search engine at 10 ppm precursor ion mass tolerance against the SwissProt Homo sapiens proteome database (20,199 entries, UniProt (http://www.uniprot.org/)). The label-free quantification and Match Between Runs were used with the following search parameters: semi-trypic digestion, fixed carbaminomethylation on cysteine, dynamic oxidation of methionine, deamidation on asparagine and glutamine, protein N-terminal acetylation with biotin labels (+ 226) of lysine residue. Less than 1% of FDR was obtained for unique labeled peptide and as well as unique labeled protein.

#### Mass spec data analysis for biotinylated peptides tagged with TMT

Raw LC-MS/MS data were processed with MaxQuant 2.7.5.0) using the Andromeda search engine. Spectra were searched against the UniProt Homo sapiens reference proteome (20,199 entries, UniProt (http://www.uniprot.org/)) appended with a common contaminant database. The enzyme specificity was Trypsin/P (C-terminal to K/R; cleavage allowed before P), permitting up to two missed cleavages. Carbamidomethyl-C was specified as a fixed modification. Oxidation-M, deamidation-N or Q, protein N-terminal acetylation or -K, biotinylation-K were set as variable modifications. The minimum peptide length was 7 aa. First-search precursor tolerance was 20 ppm and main-search tolerance 4.5 ppm; fragment tolerance was 0.02 Da for Orbitrap MS/MS FDR was controlled at 1% at the PSM, peptide, and protein levels using a target-decoy approach. The second peptides feature was enabled. Match between runs was enabled (matching time window 0.7-1.0-minute, alignment window 15-20-minute) to transfer identifications across fractions/replicates acquired under equivalent conditions. Isobaric labeling and reporter extraction. Isobaric TMT quantification was configured under Isobaric labels. The appropriate kit (TMT10) was selected, and lot-specific isotopic impurity correction factors from the manufacturer were entered via the Edit dialog. MS3 TMT (Orbitrap Eclipse): Reporter ion mode = MS3; reporter mass tolerance = 0.003–0.007 m/z; PIF filtering was applied more permissively (≥ 0.5). Protein groups were assembled using razor+unique peptides. Reverse hits, proteins only identified by site, and potential contaminants were removed prior to quantification. Unless otherwise stated, reporter intensities were taken from reporterIntensityCorrected columns (impurity-corrected), and MaxQuant’s internal global scaling to the reference channel was used across runs. For downstream analysis, protein-level reporter intensities were log_2_-transformed; proteins were retained if they had sufficient observations across conditions with Min ratio count ≥ 2 (unless noted otherwise). The full LC-MS acquisition parameters (column, gradient, resolution settings, AGC targets, max injection times, isolation windows, dynamic exclusion.

#### Mass spec data analysis for label-free quantification of biotinylated peptides (TimsTOF)

The .d files acquired in data-dependent and data-independent mode were analyzed using Spectronaut v 19.1 (Biognosys AG, Schlieren, Switzerland) with the directDIA+ feature against the Uniprot Homo sapiens protein database. Proteolysis with Trypsin was assumed to be specific, allowing for N-ragged cleavage with up to two missed cleavage sites, with a PSM FDR of 1%. Data filtering was based on the Q Value, and global normalization was utilized. Cysteine modified with iodoacetamide was set as a fixed modification. Variable modifications included biotinylation of lysine (mass shift +226 Da), oxidation of methionine, deamidation of asparagine and glutamine, deoxidation of methionine and tryptophan, and phosphorylation of serine, threonine, and tyrosine, acetylation of N-terminal and lysine, and pyro-glutamic acid conversion from glutamine and glutamic acid.

#### Data normalization and missing value imputation

Protein-level intensity values exported from MaxQuant were first filtered to remove reverse hits, potential contaminants, and proteins identified only by site. Intensities were log_2_-transformed and uploaded to NormalyzerDE (web interface, https://normalyzerde.immunoprot.lth.se/) for evaluation of normalization strategies. Based on the diagnostic reports (pooled coeffient of variation, MA plots, and PCA clustering), quantile normalization was selected as the optimal method and applied to all samples. Quantile-normalized log_2_ intensities were then exported and used for downstream statistical analysis.

For imputation of missing value, Perseus (Version 2.1.6.0, software for MaxQuant-based proteomic analysis; https://maxquant.net/perseus/) was used. The remaining missing values, assumed to arise mainly from low-abundance proteins below the detection limit, were imputed using a left-censored normal distribution. For each sample, missing entries were replaced by random values drawn from a normal distribution-shifted 1.8 standard deviations below the mean of the observed log_2_ intensities with a width of 0.3, thereby simulating low but non-zero intensities close to the detection threshold. The resulting normalized and imputed log_2_ intensities were used for all subsequent analysis.

#### Bioinformatic analysis

Perseus (Version 2.1.6.0, https://maxquant.net/perseus/) was used for analysis and visualization. The number of biological replicates for each condition is shown in Table S2 and S3.

Pairwise Pearson correlations across replicates: To assess reproducibility between biological replicates, Pairwise Pearson correlation coefficients were calculated with Perseus software. Quantile-normalized log_2_ protein intensities (exported from NormaluzerDE) before missing-value imputation were imported into Perseus, and for each pair of samples only proteins with valid intensity values in both samples were included. The resulting correlation coefficients were visualized as a clustered correlation matrix and used as a quality control metric for replicate similarity.

PCA: To visualize overall similarity between samples and major sources of variance, PCA was performed with Perceus. The filtered, quantile-normalized log_2_ protein intensity matrix with missing values imputed by left-censored normal distribution was imported into Perceus, and proteins were required to have at least one valid value across the dataset. Intensities were mean-centered by protein, and PCA was carried out using the built-in “Principal Component Analysis” function with default setting (based on the covariance matrix). Scores for principal component 1, 2 were plotted to display clustering of biological replicates and separation of experimental conditions, and the percentage of total variance explained by each principal component was reported in the corresponding plots.

GO analysis: Protein lists significantly enriched in one genotype against the other as indicated in Figure 4 and S5A, or as shown in Figure 2B, 3A, 5, 6, and S5H, were uploaded to g:profiler (https://biit.cs.ut.ee/gprofiler/gost) or Enrichr (https://maayanlab.cloud/Enrichr/) and functional enrichment of each list was performed.

To visualize Bait-Bait similarity heatmap (Figure 2B), Prey-Prey analysis (Figure 3A), and dot plots (Figure 1E, 5F), we performed SAINT analysis using SAINT implemented on a Galaxy server ^(^https://galaxy-main.usegalaxy.org/). Bait-prey matrix generated from normalized log_2_ intensities were uploaded to the Galaxy, and SAINT (SAINTexpress) was run with default parameters. And SAINT output tables generated on the Galaxy server were imported into the ProHits-Web online interface (https://prohits-viz.lunenfeld.ca/) for correlation-based visualization of the interactomes such as Bait-Bait similarity heatmap and Prey-prey analysis. SAINT output was imported into the ProHits-Web server for Bait-Bait correlation analysis. Bait-Bait similarity was quantified using the Jaccard distance score (0-1, intersection/union of SAINT-significant preys between each pair of baits), with all other parameters kept at default settings. The resulting Jaccard similarity matrix was visualized as a clustered heat map providing an overview of proximal proteomes between different types of Miro1 baits in different cell types. Each bait was manually profiled in g:profiler (https://biit.cs.ut.ee/gprofiler/gost). The same SAINT output matrix was uploaded to the ProHits-Web server for visualization of group of preys co-enriched across baits. For this analysis, quantified values (normalized log_2_ intensities) for each prey across all baits were used. Data were row-wise standardized and Pairwise Pearson correlation coefficients between preys were calculated using the default setting on the server. The resulting Prey-Prey correlation matrix was hierarchically clustered and displayed as a heatmap, and clusters of highly correlated preys were used to define functionally coherent protein groups for downstream analysis and visualization. To visualize interaction pattern for selected groups of proteins (Figure 1E and 5F), we generated Bait-Prey dot plots using the Dot Plot module in Prohits-Viz. SAINT output was uploaded into the Dot Plot Tool using the default setting, with baits displayed as columns and preys as rows. The resulting dot plots were used to illustrate differential enrichment patterns of selected protein groups across the baits.

Data analysis and visualization using Cytoscape (Figure 3E): Network-based functional visualization of the proteomic data was performed using ClueGO App (ver. 2.0.6) under Cytoscape plugin (Ver. 3.10.4; https://cytoscape.org/). Protein lists from Cluster A1 and Cluster A6 were uploaded onto ClueGO in the Cytoscape. Enrichment was tested against Gene Ontology Biological Process (GO:BP) and Cellular Component (GO:CC), using a right-sided hypergeometric test with Benjamini Hochberg correlation for multiple testing. ClueGO grouped functionally related GO terms into networks based on kappa score (kappa=0.4, default setting). The nodes represent significantly enriched GO terms/pathways. Each color denotes a specific functional group assigned by ClueGO; terms sharing ≥50% of their genes are automatically merged and colored alike. Node sizes are proportional to the -log_10_ adjusted p values (larger is more significant). The network is laid out with the organic algorithm and grouped using kappa statistics. Figure S4C: To identify proteins specifically enriched in GBM cells from whole-cell proteomics, we compared normalized protein intensities between GBM and the two types of non-GBM cell lines. For each protein, Log_2_FC was shown in x and y axis, calculated with normalized intensities. Proteins were classified as “GBM-specific” if Log_2_FC>1 in each comparison.

Figure S4D: Pathway analysis of the sorted GBM-specific proteomes as shown in (C) from Panther (https://www.pantherdb.org/tools/compareToRefList.jsp).

Figure S4E: Tissue-specific aging signatures of the sorted 120 proteomes from GTEx portal (https://www.gtexportal.org/home/).

Figure S5C-F, S7: Protein lists were first assigned to each group as described in Figure 4 and 7. The normalized and log_2_-transformed protein intensity matrix was imported into Perseus software. Profile plot visualization was plotted with the built-in profile plot function in Perseus, with individual proteins shown as thin lines. Protein lists with different cellular localizations annotated by MitoCarta3.0 and native organelle IP were individually imported for each profile plot.

Figure 7E: Functional category was analyzed by Proteomaps (https://www.proteomaps.net/) for elevated proteins upon MR3 treatment. Annotations are based on Kyoto Encyclopedia of Genes and Genomes pathways. The size of each polygon correlates with the log_2_FC(+/− MR3).

Mitochondria-associated translation set (MATS): all genes with log₂(ribosome enrichment)>1.0 in both replicates, corresponding to a relaxed threshold compared to the translatome (≥1.5) in LOCL-TL ^34^.

MitoSurf and HCPS: From total 1901 prey proteins identified from proximity labeling proteomics, we manually removed 63 of exclusively identified preys from LOV-Tb-EGFP bait. From the remaining 1838 proteins (MitoSurf), we computed a High-Confidence Proximity Score (HCPS) for each prey protein by integrating 5 rubric factors to quantify the confidence of proteins identified: (1) “E”nrichment vs. EGFP: bait-specific enrichment over a matched control bait (EGFP), (2) “S”tatistical significance (p values), (3) replicate “C”onsistency (frequency of identified peptide), (4) “B”ackground frequency (B+L-) as background correction, and (5) “subcellular “L”ocalization prior (“OMM evidence” from MitoCarta3.0/organelle IP/Uniprot CC). For enrichment score (E), normalized MS intensities were processed as log_2_ fold change (FC) between bait and EGFP samples and E was calculated independently for each run and normalized to 8 (99^th^ percentile of positive log_2_FC value, default). E=min(1, max(0,log_2_FC)/8). Statistical significance (S) was derived from t-tests with Benjamini-Hochberg correction, using the mean of the two run’s -log_10_ (p-value, capped at 6). S=min(1, −log_10_(p value)/8). Consistency score (C) was defined as the average of run-specific presence-weighted reproducibility terms. C=# of bait/# of replicate (3) * (1-clip(CV,0,1)). CV (bait) is coefficient of variation of bait replicates (linear scale) and clip to |0,1| to avoid over-penalty (=min(1, STDEV.P(MS intensity from all replicates)/Average(MS intensity from all replicates)). To strongly penalize nonspecific labeling, a nonlinear background penalty (B) was computed from 3 negative control samples (Biotin+ Light -). For each run, the detection frequency in negative controls (NC) was estimated and transformed as: B=1-(# of detection in NC/3). A localization prior (L) used prior knowledge of preys at the OMM from MitoCarta3.0, Uniprot, and mitochondrial evidence from organelle IP. 1: found in 2-3 datasets, 0.5: found only in 1 dataset, and 0: not found. The final score was calculated as: HCPS = 0.35E + 0.20S + 0.15C + 0.25B + 0.05L Proteins with > 0.6 were considered highconfidence, 0.3 - 0.6 as mediumconfidence, and < 0.3 as low-confidence. Our HCPS framework was implemented as a custom composite confidence score, conceptually grounded in established AP-MS/PL scoring approaches ^54–57^ that integrate enrichment/abundance, reproducibility, control/background frequency (MiST, CompPASS), and control-driven statistical scoring (SAINT), with contaminant frequency assessment guided by CRAPome.

#### RiboLOOM

Light-induced proximity biotinylation of polysome complexes: Cells stably expressing LOV-Tb-Miro1-WT/-KK/-OMM were seeded on 100 pi culture dishes and grown to ∼80% confluence in complete DMEM culture medium in a 37°C, CO_2_ incubator. For light-induced biotinylation, cultures were pre-equilibrated in complete medium containing 50 μM biotin for 30 minutes at 37°C. Dishes were then illuminated from below using a blue LED array indicated as above. Immediately after labeling, cells were placed on ice, washed twice with ice-cold PBS supplement with 100 μg/ml cycloheximide (CHX) to freeze translating ribosome, and processed for polysome-preserving lysis.

Polysome-preserving lysis: For all conditions, cells from one 100 pi dish were scraped in 7 ml ice-cold PBS containing 100 μg/ml CHX, collected by centrifugation (500 xg, 5 minutes, 4°C), and the supernatant was discarded. Pellets were resuspended in 600 μl of polysome lysis buffer per dish (20 mM Tris-HCl, pH 7.4, 150 mM KCl, 5 mM MgCl_2_, 1% NP-40, 1mM DTT, 100 μg/ml CHX, 40 U/ml RNase inhibitor, Protease inhibitor cocktail (EDTA-free), 0.1 U/μl TURBO DNase). Cell suspensions were incubated at 4°C for 10 minutes with occasional gentle flicking, followed by mechanical shearing through a 25 G needle (5-10 strokes) to complete lysis and release polysomes. Lysates were clarified by centrifugation at 17,000 xg for 10 minutes at 4°C, and the resulting supernatant (polysome-preserving clarified lysate) was used for streptavidin enrichment. A small aliquot (10 μl) was saved as input for normalization to bait expression level (Figure S6B).

Enrichment of biotinylated polysome-RNA complexes: Biotinylated polysome-RNA complexes were captured using Pierce streptavidin magnetic beads (88817, Thermo). Beads (50 μl) per sample were equilibrated by washing 3 times with polysome-preserving lysis buffer and resuspended at ∼10 μl settled beads. Equal amounts (600 μl) of clarified lysate from Miro1-WT, KK, and OMM samples were incubated with the pre-equilibrated beads in low-binding tubes at 4°C with end-over-end rotation for 1 hour. After incubation, tubes were placed on a magnetic rack and the post-binding supernatant was collected as the Unbound (U) fraction. Beads (Bound/B fraction) were washed 4 times with high-stringency wash buffer (20 mM Tris-HCl, pH 7.4, 300 mM KCl, 5 mM MgCl_2_, 1% NP-40, 1 mM DTT, 100 μg/ml CHX, RNase inhibitor and protease inhibitor cocktail). Each wash consisted of gentle resuspension, 5-minute rotation at 4°C, and magnetic separation. A final wash in low salt buffer (20 mM Tris-HCl, pH 7.4, 150 mM KCl, 5 mM MgCl_2_, 1% NP-40) was performed to remove excess salt and detergent prior to RNA extraction. RNA extraction from B and U: After the last wash, beads were resuspended directly in 200 μl proteinase K buffer (50 mM Tris-HCl, pH 7.5, 10 mM EDTA, 1% SDS) supplemented with proteinase K (0.2 mg/ml). Bead suspensions were incubated at 55°C for 30 minutes occasional gentle mixing to digest bead-bound proteins and release polysome-RNA complexes. Following proteolysis, the post-binding supernatant from the beads was collected in new tubes and 200 μl of TRIzol reagent was added to the beads. The TRIzol-bead suspensions incubated for an additional 20 minutes at 37°C to ensure complete denaturation of proteins and dissociation of ribonucleoprotein complexes. The content was then combined with the previous proteinase K-treated supernatant. Chloroform was added at 0.2 volumes relative to TRIzol, samples were vigorously vortexed for 15 seconds, and incubated at room temperature for 3 minutes, and centrifuged at 12,000 xg for 10 minutes at 4°C to separate phase. The upper aqueous phase was transferred to a fresh tube and extracted once more with an equal volume of acid phenol-chloroform (pH 4.5). After vortexing and centrifugation (12,000 xg, 10 minutes, 4°C), the final aqueous phase was collected. RNA was precipitated by adding 2.5 volumes of isopropanol with 1 μl of glycogen as carrier. Samples were centrifuged at 16,000 xg for 10 minutes at 4°C, and the pellets were washed with 70% ethanol. After a brief air-dry, RNA pellets were resuspended in 20 μl RNase-free water and stored at -80°C until reverse transcription. The U was processed in parallel using the upper workflow for RNA extraction. TRIzol–chloroform phase separation, subsequent acid phenol extraction, isopropanol precipitation, washing with ethanol, and resuspension in 100 μl of RNase-free water was performed exactly as described for the Bound fraction, ensuring that Bound and Unbound RNAs were isolated using identical conditions. RNA concentration and purity (A260/280 ratio) for both fractions were determined by Nano drop, and equal RNA inputs (1 μg) were used in downstream reverse transcription and RT-qPCR.

Reverse transcription: Typically, 1 μg RNA from B and U was used for cDNA synthesis according to the manufacturer’s instructions (Abclonal) in a 20 μl reaction using a reverse transcription mastermix (5x, RK20428, Abclonal): 65°C primer annealing, 55°C extension, 70°C inactivation). Before qPCR, cDNA was typically diluted to 1:2 in nuclease-free water.

Quantitative PCR: About 20 ng cDNA estimated based on RNA amount per qPCR reaction was used to keep Ct values within dynamic range. Gene-specific primers were designed to amplify 80-150 bp products with melting temperature of 60°C, verified in Origene (https://www.origene.com/), and synthesized from ELIM biopharm (https://www.elimbio.com). Reactions were set up in 96-well plates using a SYBR Green-based master mix (2x, RK21205, Abclonal) in a final volume of 20 μl according to the manufacturer’s instructions (Abclonal). qPCR was performed on a real-time instrument (Bio-Rad) with the following cycling program:

1. 95°C for 3 minutes (initial denaturation)
2. 40 cycles of: 95°C for 10 seconds and 60°C for 30 seconds (annealing/extension, with fluorescence acquisition)
3. Melt curve analysis to verify single specific products

Each gene was measured in biological triplicates for both Bound and Unbound fractions from each condition (Miro1-WT, OMM, KK).

Lt (local translation index) ^58^: B: bound, U: unbound

Lt=log_2_(B/U)=−(ΔCqB−ΔCqU)

Lt = log_2_(2^-Cq,B^/2^-Cq,U^).

Lt = 1: 2-fold enrichment in Bound relative to Unbound; Lt=0: no enrichment in either Bound or Unbound; L = -2: 4-time enrichment in Unbound. Cutoff for no enrichment: Lt (average)<0.2.

#### AHA assay

HEK293T cells stably expressing dCas9-KRAB or dCas9-KRAB with sgRHOT1 were cultured in DMEM supplemented with 10% FBS and penicillin-streptomycin in a 37°C, 5% CO_2_ incubator. Myc-Miro1-WT/KK was transfected into Miro1-KD cells through PEI-based transfection as previously described. The next day after the transfection, cells at ∼80% confluency were washed once with pre-warm PBS and pre-incubated for 1 hour in methionine-free DMEM (21013024, Gibco) supplemented with 10% dialyzed FBS, 0.2 mM L-cysteine (AAJ6374514, Fisher), and 25 mM HEPES (pH 7.4) for metabolic labeling of newly synthesized proteins. L-Azidohomoalanine (AHA, C10102, Invitrogen) was then added to a final concentration 50 μM and cells were pulse-labeled for 15 minutes at 37°C. After the pulse, cells were placed on ice, washed twice with ice-cold PBS, and collected by scraping into ice-cold mitochondrial isolation buffer (MIB; 250 mM sucrose, 10 mM HEPES pH 7.4, 1mM EDTA, protease inhibitor cocktail). Cell suspensions were homogenized on ice with a pestle and needle (20-30 strokes) and cleared at 600 xg for 5 minutes at 4°C to remove nuclei and debris. The post-nuclear supernatant was centrifuged at 10,000 xg for 10 minutes at 4°C to pellet crude mitochondria. The resulting supernatant was collected as the cytosolic fraction and the mitochondrial pellet was washed once in MIB (10,000 xg, 10 minutes, 4°C) and gently resuspended in lysis buffer. Protein concentrations in mitochondrial and cytosolic fractions were determined by BCA assay. To validate the purity of each fraction prior to AHA-labeling, a small aliquot (20 μl) of each fraction taken before Click-iT chemistry was analyzed by SDS-PAGE. For Click-iT conjugation of AHA-labeled proteins, 200 μg protein from each fraction was adjusted to 200 μl with IP lysis buffer (25 mM, Tris-HCl pH 7.4, 150 mM NaCl, 1% NP-40, 1 mM EDTA, protease inhibitor cocktail). Click-iT reactions were performed using a biotin-PEG4-alkyne Click-iT chemistry system according to the manufacturer’s instructions, with a final reaction volume of 300 μl per sample. Briefly, biotin-PEG4-alkyne (B10185, Invitrogen) and Click-iT Protein Reaction Buffer kit including reaction buffer, CuSO4 solution, and reducing/ligand additives (C10276, Invitrogen) were added sequentially to the lysate, mixed thoroughly, and incubated for 20 minutes at room temperature with gentle rotation. Following Click-iT chemistry, proteins were precipitated by adding 4 volumes of methanol, 1 volume of chloroform, and 3 volumes of water (final ratio: lysate:MeOH:CHCl_3_:H_2_O = 1:4:1:3), vortexed vigorously after each addition, and centrifuged at 14,000 xg for 5 minutes at 4°C. The upper aqueous phase was carefully removed without disturbing the interphase protein disk. The lower organic phase was then aspirated, leaving the white/opaque protein disk at the interface. The protein pellet was washed twice with 500 μl cold MeOH (Vortex, 14,000 xg, 5 minutes, 4°C), air-dried briefly until residual solvent evaporated, and resuspended in 100 μl IP lysis buffer. Pellets were solubilized by repeated pipetting and vortexing at room temperature for 10 minutes, followed by centrifugation at 14,000 xg for 5 minutes at 4°C to remove insoluble debris. The cleared supernatant was used for streptavidin enrichment. It was incubated with 20 μl packed streptavidin magnetic beads (pre-washed three times with IP lysis buffer) for 1 hour at 4°C on a rotator. Beads were washed 5 times with 200 μl ice-cold IP lysis buffer and once with PBS. Bound proteins were eluted by boiling beads in 20 μl of 1X SDS-PAGE sample buffer containing 50 mM DTT at 95°C for 10 minutes. Eluates were resolved by SDS-PAGE and transferred to nitrocellulose membranes. Global AHA incorporation and enrichment were assessed by probing membranes with IR800CW conjugated streptavidin as described above. For analysis of specific nascent target hits, membranes were blocked in 5% skimmed milk in TBST and incubated with primary and secondary antibodies described above.

#### Website building

All datasets analyzed in this work are accessible through our laboratory’s interactive web resource, MiroScape (https://miroscape.github.io/MiroScape). Data relevant to this study are available under the “MitoSurf” module and “MiroProteome” → “Glioma and PD Cells” tab. The MitoSurf dataset provides LOV-Turbo-Miro1 proximity labeling data across contexts. Mean MS values are shown and expressed as heatmaps. The MiroProteome-Glioma and PD Cells dataset contains proteomic data from glioma cells and iDAs (WT and SNCA-A53T), with and without MR3 treatment. Users can search for proteins by gene names and view their average MS intensity results and heatmap visualizations in our data. The website is hosted on GitHub Pages, using the React framework for front-end development and react-plotly.js for data visualization. All source code is openly available at: https://github.com/miroscape/MiroScape.git. Software and algorithms: React version 9.2.0, Meta, https://react.dev/; Ant Design version 5.27.4, Ant Group, https://ant.design/; react-router-dom version 7.9.4, Remix, https://reactrouter.com/, iframe-resizer version 5.1.5, Bradshaw, https://github.com/davidjbradshaw/iframe-resizer; react-plotly.js version 2.6.0, Plotly, https://plotly.com/javascript/.

#### Database used for analysis

MitoCarta3.0 ^7^, native organelle IP database ^5,10^, and the Human Protein Atlas (HPA) subcellular localization data (https://www.proteinatlas.org/about/download, version 24.0) were used.

#### Visualization of ribosomal proteomes

The cryo-EM structure of the human mitochondrial ribosome (PDB: 3J9M), cytosolic ribosome (PDB: 8JDM, 8PJ1, 9G8M, 8XXN), prohibitin complex (PDB: 9O6S), two types of Human tRNA synthetases predicted from AF3 (AF-Q9NSE4) and cytosolic histidine tRNA synthetase (PDB: 4X5O), TOM complexes (PDB: 7CP9, 8XVA, 7E4I), and AF3-predicted metaxin complexes was loaded with biotin-labeled sites, individually. The structure was rendered in Pymol.

### QUANTIFICATION AND STATISTICAL ANALYSIS

Throughout the paper, the distribution of data points is expressed as box-whisker or dot plot, except otherwise stated. For box-whisker, the whiskers are min to max. One-Way ANOVA was performed for comparing multiple groups. Statistical analyses were performed using the Prism software or Excel. For all experiments, between 3 and 6 biological samples were used or independent experiments were performed. The number and meaning of experimental replications (n) can be found in corresponding figure legends. No statistical methods were used to predetermine sample sizes, but the experiments, and biological replicates were determined based on the nature of the experiments (since it is often difficult to assess an outcome that follows a normal distribution in our experiments), degree of variations, and published papers describing similar experiments. Precise p values are labeled in figures.

