## Supplemental Figures and Legends for "Optogenetic Proximity Labeling Maps Spatially Resolved Mitochondrial Surface Proteomes and a Locally Regulated Ribosome Pool"

**Supplemental Materials**

**Supplemental Figure Legends**

**Figure S1 Validation of the constructs.**

(A) Confocal microscopy images show colocalization of V5-tagged baits and the mitochondrial marker (MitoTracker Deep Red) or peroxisome marker (EGFP-SRL) in GBM cells. Scale bar: 25 μm. The lower row shows images of Pearson's correlation between V5 and MitoTracker Deep Red (mtDR) or SRL, generated from the same image above by Image J.

(B) Live imaging images of TMRM and MitoTracker Green (independent of membrane potential) in HEK293T cells. Scale bar: 25 μm. Quantification of the mean intensity of TMRM/MitoTracker Green within one cell, averaged across 5 cells per image. One-Way Anova PostHoc Tukey Test. n.s.: not significant. n=4.

(C) Western blots of biotinylated proteins from each stable cell line. Similar results were seen more than 3 times.

**Figure S2 Subcellular localization of biotinylated proteins.**

Confocal microscopy images show bait (V5 or EGFP) and biotinylated protein (Streptavidin) localization, with or without light in HEK293T and GBM cells (A), iPSCs (B), and iNs (C). Scale bars: 25 μm.

**Figure S3 Workflow and overview of our PL data.**

(A) Schematic illustration of our PL proteomic strategy.

(B) MS2/MS3 spectrum of TMT-labeled biotin-modified peptides corresponding endogenous TRAK1 from DDA label quantification.

(C) MS2 spectrum of biotin-modified peptides corresponding to endogenous TRAK1 from DIA label-free quantification.

(D) The MS intensity of identified sites (Lysine residues) of TRAK1 in HEK293T and GBM cells.

(E) PCA analysis of our cell-type-specific PL proteomics. PCA was performed on $log_2$-transformed, medium-centered, and imputed MS intensity of all quantified proteins under Perseus plugin. x-axis (PC1) explains 40.3% of the total variance. y-axis (PC2) explains 11.3% of the total variance. Color code indicates cell type and bait type.

**Figure S4 Workflow and overview of quantitative whole-cell proteomics.**

(A) Schematic illustration of whole-cell proteomic strategy.

(B) PCA analysis of our cell-type-specific quantitative whole-cell proteomics. PCA was performed on $log_2$-transformed, medium-centered, and imputed MS intensity of all quantified proteins under Perseus plugin. x-axis (PC1) explains 61.2% of the total variance. y-axis (PC2) explains 16.4% of the total variance. Color code indicates cell type and bait type.

(C) The sorted GBM-specific proteomes from the whole-cell proteomic data. Scatter plot shows every quantified protein by its dual $log_2FC$. x-axis is $log_2FC$ (GBM/HEK293T) and y-axis is $log_2FC$ (GBM/iDA) under the basal condition. Grey color marks proteins whose abundance does not exceed the enrichment cut-off ($Log_2FC>1$) in either comparison. Red-colored dots highlight the GBM-specific protein set (120 proteins).

(D) Pathway analysis of the sorted GBM-specific proteins shown in (C) from Panther. Blue box indicates specific disease signatures.

(E) Tissue-specific aging signatures of the sorted 120 proteins from GTEx portal. The sorted proteins are significantly upregulated in the aged brain.

(F) Cell-type-specific protein abundance of representative proteins whose signals are changed upon MR3 treatment in the PL data.

**Figure S5 Miro1 proximally interacts with multiple organelles and translation machinery.**

(A-B) GO analysis of Biological Process of Miro1-OMM (A) or Miro1-WT-enriched preys (B) (from Figure 4G and H) in HEK (blue) and GBM cells (red). GO terms are significantly enriched in the dataset generated from Enrichr (https://maayanlab.cloud/Enrichr/).

(C-G) The profile plot ($log_2$-normalized intensity traces for each protein) and subcellular annotation of the selected proteomic components of Miro1-OMM in HEK (C), Miro1-OMM in GBM (D), Miro1-OMM in iN (E), Miro1-WT in HEK (F), and Miro1-WT in GBM (G), described in Figure 4. Each proteomic component was manually classified from public proteomic databases (Mitocarta3.0 and native organelle IP) and the corresponding representative proteomes are shown.

(H) GO term analysis of short and long CDS groups classified from 183 filtered MitoCarta3.0 preys generated with Enrichr.

(I) Workflow and results of comparing our MitoSurf preys with MATS.

**Figure S6 Miro1-KK-specific proteomes and functions.**

(A) The dual-comparison proteome map by a correlation scatter plot. Each point is a protein with by the $log_2FC$ measured for Miro1-WT versus Miro1-KK in two cell models. x axis is $log_2FC$ (GBM: WT/KK) and y-axis is $log_2FC$ (HEK: WT/KK). Positive values indicate higher levels in the WT samples and negative values indicate enrichment in KK. Proteins are partitioned into eight clustered (Group I-VIII) on the basis of their paired fold change pattern: $\geq$1.5 or $\leq$-1.5. Colors of the protein labels match the group colors and grey-colored proteins are not selected: Group I (brick-red, 14 proteins, enriched in Miro1-WT in both cell types), Group II (steel-blue, 255 proteins, enriched in GBM, Miro1-WT), Group III (turquoise, 33 proteins, enriched in HEK, Miro1-WT), Group IV (sage-green, 11 proteins, enriched in Miro1-KK in both cell types), Group V (orchid-magenta, 287 proteins, enriched in GBM, Miro1-KK), Group VI (amber, 73 proteins, enriched in HEK, Miro1-KK), Group VII (mustard-yellow, 10 proteins, enriched in GBM, Miro1-WT and HEK, Miro1-KK), Group VIII (olive-green, 5 proteins, enriched in GBM, Miro1-KK and HEK, Miro1-WT).

(B) Bait expression (Miro1-WT, -KK, and -OMM) immunoblot from HEK293T cells exposed to biotin and blue light for normalizing RNA pools captured from polysomes. Red arrows indicate the major V5-positive signal. Right: V5 signal intensity (arbitrary unit) was quantified from image J. n=3.

(C-D) RNAs shown in Figure 6F are normalized to the total intensity of cytoplasmic ribosome subunits (C) or AKAP1(D) captured by each bait from the PL proteomics.

(E) Vectors used in silencing *Miro1* with CRISPRi under the bidirectional UbC/U6 promoter cassette. The UbC promoter drives expression of 3x FLAG-dCas9-KRAB-T2A-EGFP, while the opposing U6 promoter drives either a non-targeting sgRNA scaffold (top) or sgRNAs matching part of the 5'-UTR of *Miro1* (bottom), enabling co-expression of fluorescent dCas9-KRAB and the indicated sgRNA from a single vector.

(F) RT-qPCR analysis of indicated genes in HEK293T cells expressing dCas9-KRAB with or without sgRHOT1. Gene expression was normalized to GAPDH and plotted as fold change relative to control (red). n=3.

(G) Immunoblots of Miro1 and ACTB (loading control) in HEK293T expressing dCas9-KRAB in the presence or absence of sgRHOT1.

(H) Immunoblots as indicated. Cells were transfected with a series of amounts of Myc-Miro1-WT or -KK (one 12-well). Ponceau S staining shows total proteins of the same gel.

(I) Similar to (H), but 0.25 ug of Myc-Miro1-WT or -KK was transfected. Right: Quantification of the anti-Miro1 bands normalized to ACTB. n=3.

(J) Representative confocal images of HEK293T cells as indicated, live stained with MitoTracker Green (membrane potential independent) and TMRM. Scale bar: 25 μm. Right: Quantification of the mean intensity of TMRM normalized to MitoTracker Green within each cell, averaged across 5 cells per image. n=4.

(K) A representative confocal image of HEK293T cells as indicated stably expressing Mito-mKeima (red, excited at 586 nm) stained with MitoTracker Deep Red (cyan), to demonstrate the correct localization of Mito-mKeima. Scale bar: 25 μm. Right: 2D intensity plot of MitoTracker Deep Red versus Mito-mKeima, generated from the confocal image with ImageJ, illustrating the tight colocalization of Mito-mKeima with mitochondria (indicated by R-squared values).

(L) Quantification of Mito-mKeima intensities (excited at 586/440 nm) using a plate reader. Right: CCCP-triggered mitophagy: The mitophagy index at baseline minus that with CCCP. n=6.

(M) Plots of OCR and ECAR are shown.

(N) Quantification of basal OCR (left) and basal ECAR (right) from the plots shown in (M). n=4.

One-Way Anova PostHoc Tukey Test (B, C, D, F, I, J, L, N).

**Figure S7 MR3-modulated Miro1 proximity interactomes.**

The profile plot ($log_2$-normalized intensity traces for each protein) and subcellular annotation of MR3-responsive Miro1-proximal interacting proteins identified from HEK and GBM cells, described in Figure 7D (Groups I-IV). Proteins are further categorized based on proteomic databases (Mitocarta3.0 and native organelle IP).

**Supplemental Tables**

**Table S1 Information of plasmids generated or used in this study and primers.**

**Table S2 Complete datasets of PL-derived proteomics.**

**Table S3 Complete datasets of whole-cell proteomics.**

**Table S4 Protein hits enriched in Miro1-WT relative to Miro1-KK in PL datasets.**

FigS1

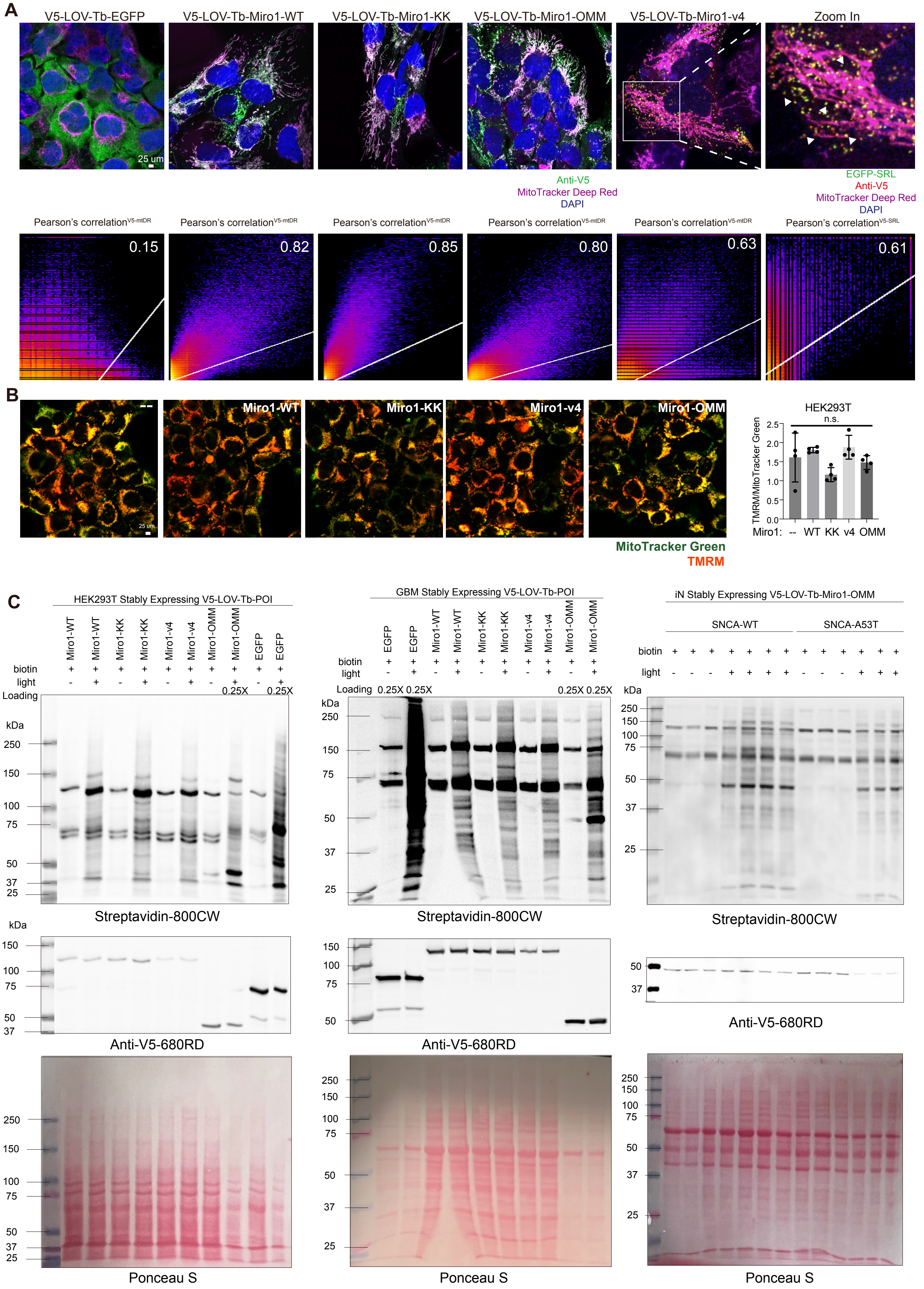
A
V5-LOV-Tb-EGFP
V5-LOV-Tb-Miro1-WT
V5-LOV-Tb-Miro1-KK
V5-LOV-Tb-Miro1-OMM
V5-LOV-Tb-Miro1-v4
Zoom In
25 um
Anti-V5
MitoTracker Deep Red
DAPI
EGFP-SRL
Anti-V5
MitoTracker Deep Red
DAPI
Pearson's correlation V5-mtDR
0.15
0.82
0.85
0.80
0.63
Pearson's correlation V5-SRL
0.61
B
--
Miro1-WT
Miro1-KK
Miro1-v4
Miro1-OMM
25 um
MitoTracker Green
TMRM
HEK293T
n.s.
TMRM/MitoTracker Green
Miro1: -- WT KK v4 OMM
C
HEK293T Stably Expressing V5-LOV-Tb-POI
Miro1-WT Miro1-WT Miro1-KK Miro1-KK Miro1-v4 Miro1-v4 Miro1-OMM Miro1-OMM EGFP EGFP
biotin
light
Loading
0.25X 0.25X
kDa
Streptavidin-800CW
Anti-V5-680RD
Ponceau S
GBM Stably Expressing V5-LOV-Tb-POI
EGFP EGFP Miro1-WT Miro1-WT Miro1-KK Miro1-KK Miro1-v4 Miro1-v4 Miro1-OMM Miro1-OMM
0.25X 0.25X 0.25X 0.25X
iN Stably Expressing V5-LOV-Tb-Miro1-OMM
SNCA-WT
SNCA-A53T

FigS2

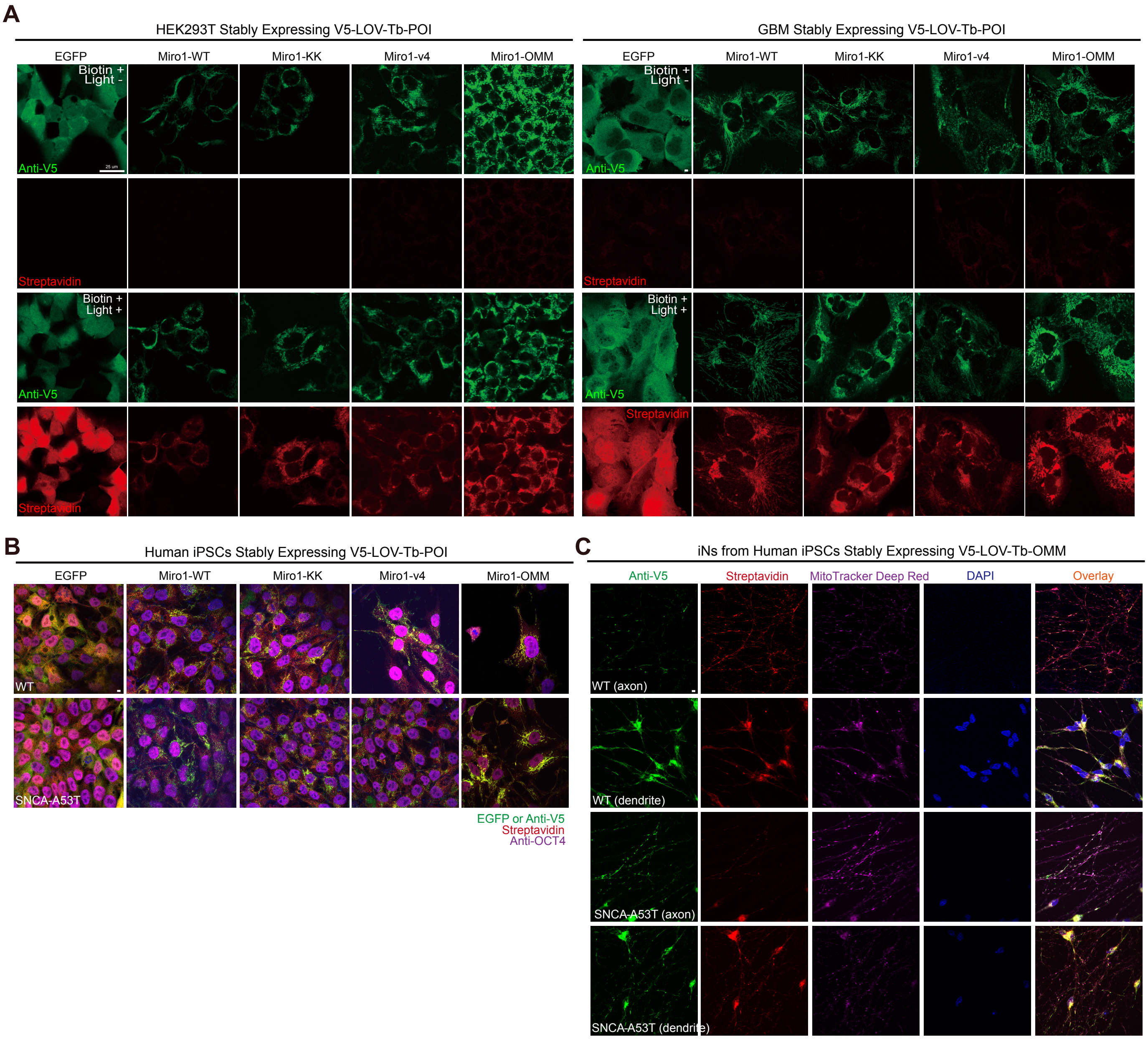
A
HEK293T Stably Expressing V5-LOV-Tb-POI
GBM Stably Expressing V5-LOV-Tb-POI
EGFP
Miro1-WT
Miro1-KK
Miro1-v4
Miro1-OMM
Biotin +
Light -
Biotin +
Light +
Anti-V5
Streptavidin
B
Human iPSCs Stably Expressing V5-LOV-Tb-POI
WT
SNCA-A53T
EGFP or Anti-V5
Streptavidin
Anti-OCT4
C
iNs from Human iPSCs Stably Expressing V5-LOV-Tb-OMM
Anti-V5
Streptavidin
MitoTracker Deep Red
DAPI
Overlay
WT (axon)
WT (dendrite)
SNCA-A53T (axon)
SNCA-A53T (dendrite)

FigS3

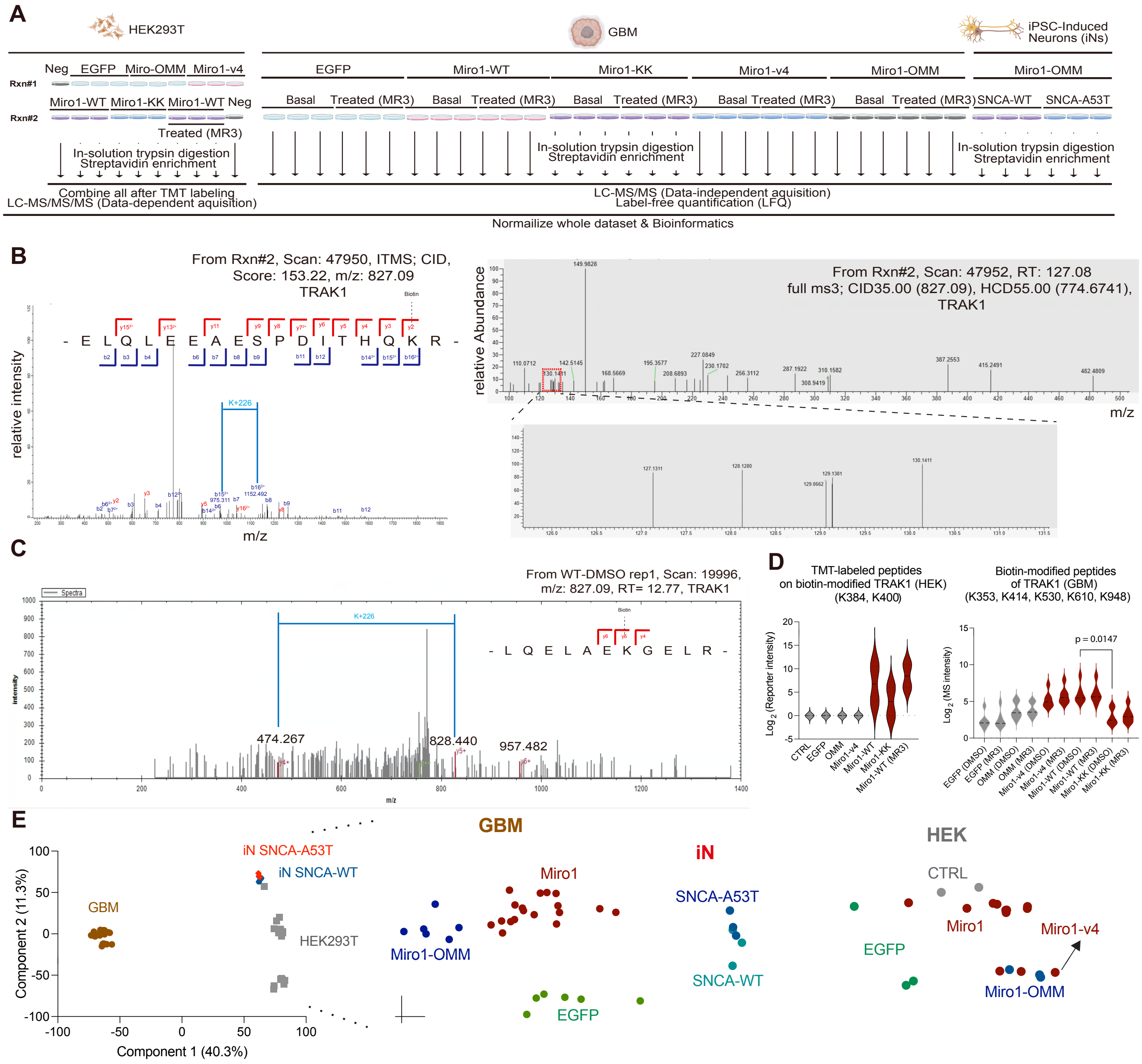

A
HEK293T
GBM
iPSC-Induced Neurons (iNs)
Neg EGFP Miro-OMM Miro1-v4
Rxn#1
Miro1-WT Miro1-KK Miro1-WT Neg
Rxn#2
Treated (MR3)
In-solution trypsin digestion
Streptavidin enrichment
Combine all after TMT labeling
LC-MS/MS/MS (Data-dependent aquisition)
EGFP
Miro1-WT
Miro1-KK
Miro1-v4
Miro1-OMM
Basal Treated (MR3)
LC-MS/MS (Data-independent aquisition)
Label-free quantification (LFQ)
SNCA-WT SNCA-A53T
Normailize whole dataset & Bioinformatics
B
From Rxn#2, Scan: 47950, ITMS; CID, Score: 153.22, m/z: 827.09
TRAK1
relative intensity
m/z
- E L Q L E E A E S P D I T H Q K R -
K+226
From Rxn#2, Scan: 47952, RT: 127.08
full ms3; CID35.00 (827.09), HCD55.00 (774.6741),
TRAK1
relative Abundance
m/z
C
From WT-DMSO rep1, Scan: 19996,
m/z: 827.09, RT= 12.77, TRAK1
- L Q E L A E K G E L R -
K+226
474.267
828.440
957.482
D
TMT-labeled peptides
on biotin-modified TRAK1 (HEK)
(K384, K400)
Log2 (Reporter intensity)
CTRL
EGFP
OMM
Miro1-v4
Miro1-WT
Miro1-KK
Miro1-WT (MR3)
Biotin-modified peptides
of TRAK1 (GBM)
(K353, K414, K530, K610, K948)
Log2 (MS intensity)
p = 0.0147
EGFP (DMSO)
EGFP (MR3)
OMM (DMSO)
OMM (MR3)
Miro1-v4 (DMSO)
Miro1-v4 (MR3)
Miro1-WT (DMSO)
Miro1-WT (MR3)
Miro1-KK (DMSO)
Miro1-KK (MR3)
E
Component 2 (11.3%)
Component 1 (40.3%)
GBM
iN SNCA-A53T
iN SNCA-WT
HEK293T
GBM
Miro1
Miro1-OMM
EGFP
iN
SNCA-A53T
SNCA-WT
HEK
CTRL
Miro1
Miro1-v4
EGFP
Miro1-OMM

FigS4

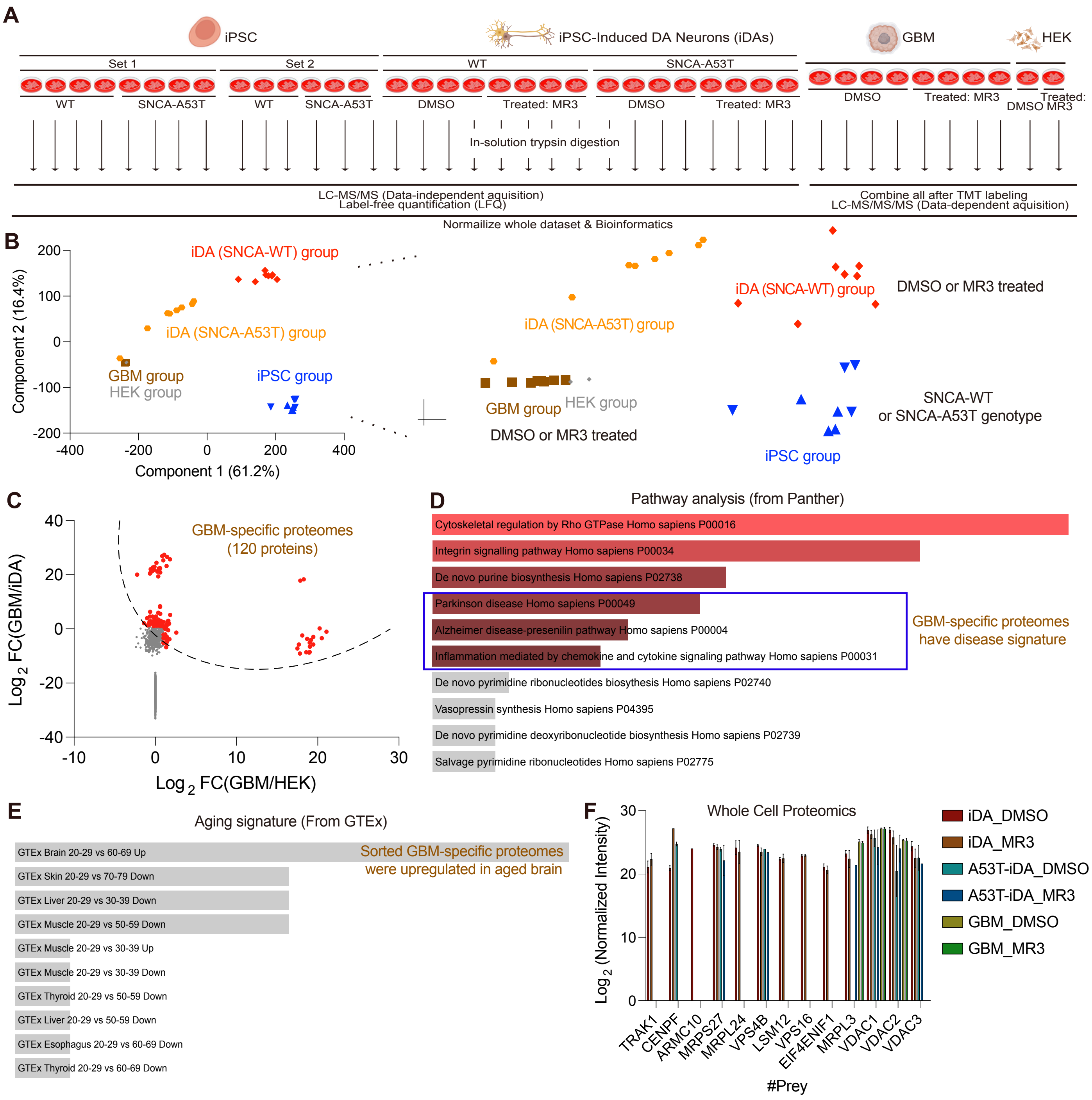
A
iPSC
iPSC-Induced DA Neurons (iDAs)
GBM
HEK
Set 1
Set 2
WT
SNCA-A53T
WT
SNCA-A53T
WT
SNCA-A53T
DMSO
Treated: MR3
DMSO
Treated: MR3
DMSO
Treated: MR3
Treated:
DMSO MR3
In-solution trypsin digestion
LC-MS/MS (Data-independent aquisition)
Label-free quantification (LFQ)
Combine all after TMT labeling
LC-MS/MS/MS (Data-dependent aquisition)
Normalize whole dataset & Bioinformatics
B
iDA (SNCA-WT) group
iDA (SNCA-A53T) group
GBM group
HEK group
iPSC group
Component 2 (16.4%)
Component 1 (61.2%)
DMSO or MR3 treated
SNCA-WT
or SNCA-A53T genotype
C
GBM-specific proteomes
(120 proteins)
Log 2 FC(GBM/iDA)
Log 2 FC(GBM/HEK)
D
Pathway analysis (from Panther)
Cytoskeletal regulation by Rho GTPase Homo sapiens P00016
Integrin signalling pathway Homo sapiens P00034
De novo purine biosynthesis Homo sapiens P02738
Parkinson disease Homo sapiens P00049
Alzheimer disease-presenilin pathway Homo sapiens P00004
Inflammation mediated by chemokine and cytokine signaling pathway Homo sapiens P00031
De novo pyrimidine ribonucleotides biosythesis Homo sapiens P02740
Vasopressin synthesis Homo sapiens P04395
De novo pyrimidine deoxyribonucleotide biosynthesis Homo sapiens P02739
Salvage pyrimidine ribonucleotides Homo sapiens P02775
GBM-specific proteomes
have disease signature
E
Aging signature (From GTEx)
GTEx Brain 20-29 vs 60-69 Up
GTEx Skin 20-29 vs 70-79 Down
GTEx Liver 20-29 vs 30-39 Down
GTEx Muscle 20-29 vs 50-59 Down
GTEx Muscle 20-29 vs 30-39 Up
GTEx Muscle 20-29 vs 30-39 Down
GTEx Thyroid 20-29 vs 50-59 Down
GTEx Liver 20-29 vs 50-59 Down
GTEx Esophagus 20-29 vs 60-69 Down
GTEx Thyroid 20-29 vs 60-69 Down
Sorted GBM-specific proteomes
were upregulated in aged brain
F
Whole Cell Proteomics
Log 2 (Normalized Intensity)
TRAK1
CENPF
ARMC10
MRPS27
MRPL24
VPS4B
LSM12
VPS16
EIF4ENIF1
MRPL3
VDAC1
VDAC2
VDAC3
#Prey
iDA_DMSO
iDA_MR3
A53T-iDA_DMSO
A53T-iDA_MR3
GBM_DMSO
GBM_MR3

FigS5

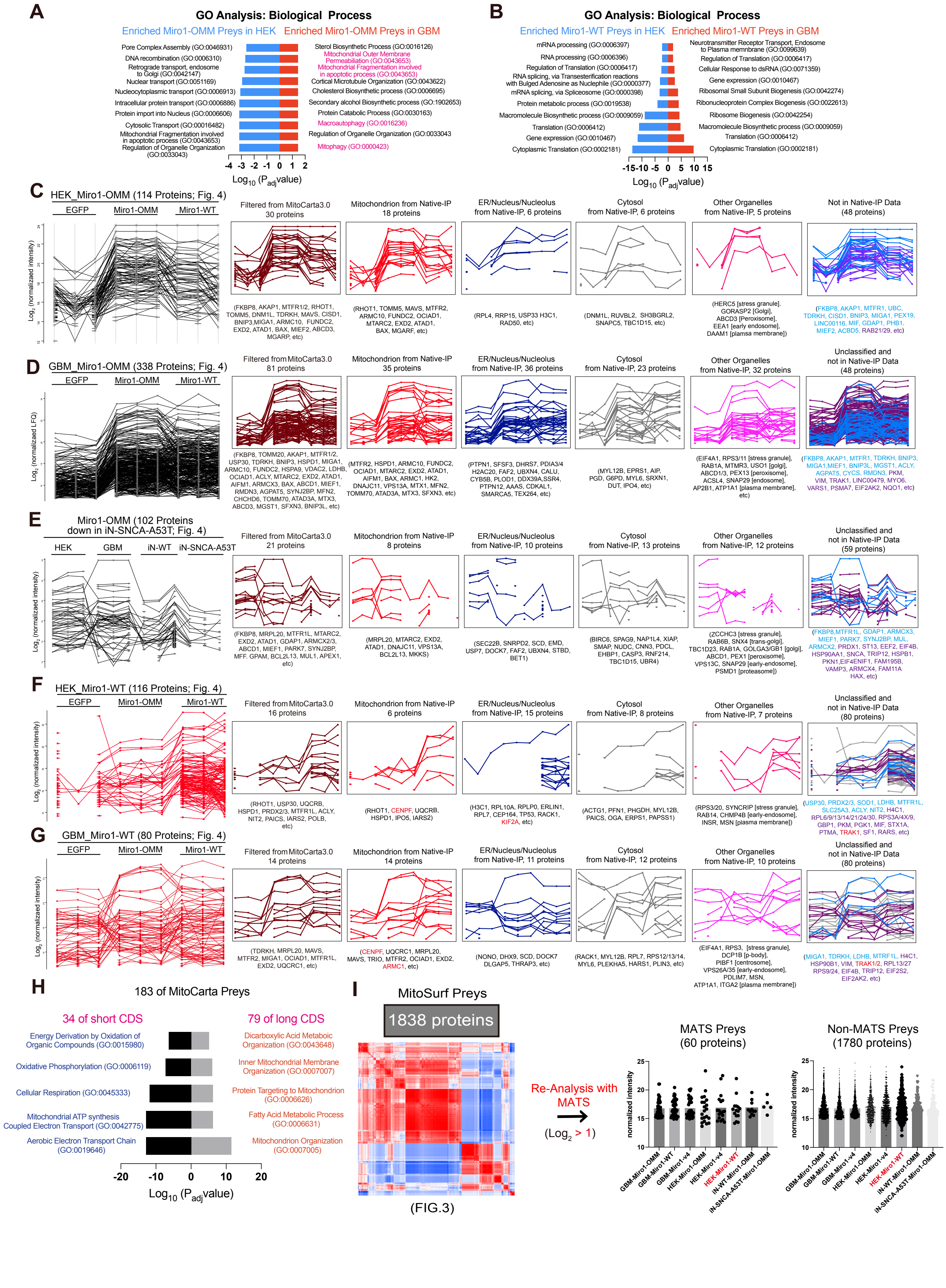
A
GO Analysis: Biological Process
Enriched Miro1-OMM Preys in HEK
Enriched Miro1-OMM Preys in GBM
B
GO Analysis: Biological Process
Enriched Miro1-WT Preys in HEK
Enriched Miro1-WT Preys in GBM
C
HEK_Miro1-OMM (114 Proteins; Fig. 4)
D
GBM_Miro1-OMM (338 Proteins; Fig. 4)
E
Miro1-OMM (102 Proteins down in iN-SNCA-A53T; Fig. 4)
F
HEK_Miro1-WT (116 Proteins; Fig. 4)
G
GBM_Miro1-WT (80 Proteins; Fig. 4)
H
183 of MitoCarta Preys
34 of short CDS
79 of long CDS
I
MitoSurf Preys
1838 proteins
(FIG.3)
Re-Analysis with MATS
MATS Preys (60 proteins)
Non-MATS Preys (1780 proteins)

FigS6

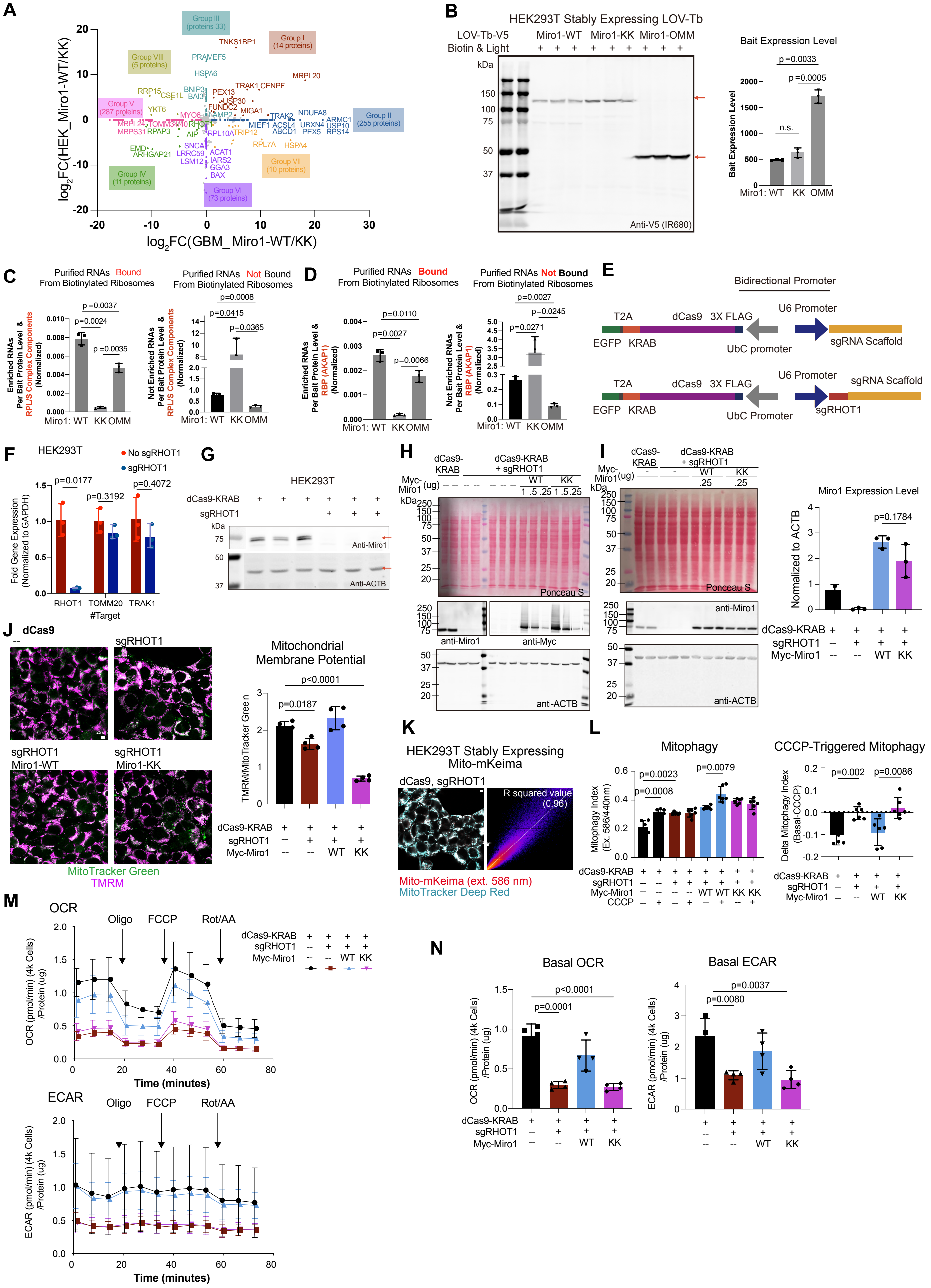
A
B
HEK293T Stably Expressing LOV-Tb
Bait Expression Level
C
Purified RNAs Bound From Biotinylated Ribosomes
Purified RNAs Not Bound From Biotinylated Ribosomes
D
E
Bidirectional Promoter
F
HEK293T
G
H
I
Miro1 Expression Level
J
Mitochondrial Membrane Potential
MitoTracker Green
TMRM
K
HEK293T Stably Expressing Mito-mKeima
Mito-mKeima (ext. 586 nm)
MitoTracker Deep Red
L
Mitophagy
CCCP-Triggered Mitophagy
M
OCR
ECAR
Time (minutes)
N
Basal OCR
Basal ECAR

FigS7

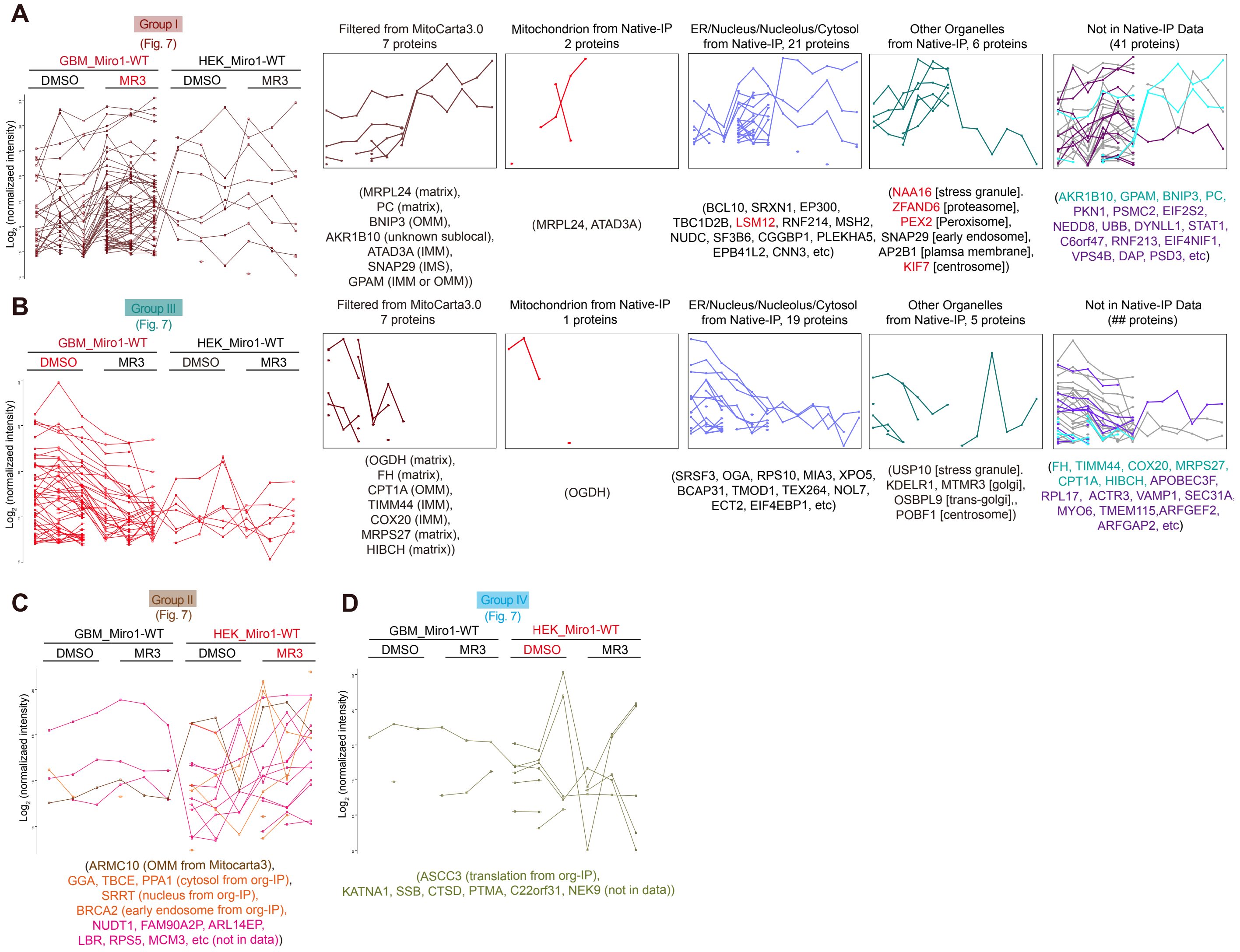

A
Group I
(Fig. 7)
GBM_Miro1-WT
HEK_Miro1-WT
DMSO
MR3
DMSO
MR3
Log2 (normalized intensity)
Filtered from MitoCarta3.0
7 proteins
Mitochondrion from Native-IP
2 proteins
ER/Nucleus/Nucleolus/Cytosol
from Native-IP, 21 proteins
Other Organelles
from Native-IP, 6 proteins
Not in Native-IP Data
(41 proteins)
(MRPL24 (matrix),
PC (matrix),
BNIP3 (OMM),
AKR1B10 (unknown sublocal),
ATAD3A (IMM),
SNAP29 (IMS),
GPAM (IMM or OMM))
(MRPL24, ATAD3A)
(BCL10, SRXN1, EP300,
TBC1D2B, LSM12, RNF214, MSH2,
NUDC, SF3B6, CGGBP1, PLEKHA5,
EPB41L2, CNN3, etc)
(NAA16 [stress granule].
ZFAND6 [proteasome],
PEX2 [Peroxisome],
SNAP29 [early endosome],
AP2B1 [plamsa membrane],
KIF7 [centrosome])
(AKR1B10, GPAM, BNIP3, PC,
PKN1, PSMC2, EIF2S2,
NEDD8, UBB, DYNLL1, STAT1,
C6orf47, RNF213, EIF4NIF1,
VPS4B, DAP, PSD3, etc)
B
Group III
(Fig. 7)
GBM_Miro1-WT
HEK_Miro1-WT
DMSO
MR3
DMSO
MR3
Log2 (normalized intensity)
Filtered from MitoCarta3.0
7 proteins
Mitochondrion from Native-IP
1 proteins
ER/Nucleus/Nucleolus/Cytosol
from Native-IP, 19 proteins
Other Organelles
from Native-IP, 5 proteins
Not in Native-IP Data
(## proteins)
(OGDH (matrix),
FH (matrix),
CPT1A (OMM),
TIMM44 (IMM),
COX20 (IMM),
MRPS27 (matrix),
HIBCH (matrix))
(OGDH)
(SRSF3, OGA, RPS10, MIA3, XPO5,
BCAP31, TMOD1, TEX264, NOL7,
ECT2, EIF4EBP1, etc)
(USP10 [stress granule].
KDELR1, MTMR3 [golgi],
OSBPL9 [trans-golgi],,
POBF1 [centrosome])
(FH, TIMM44, COX20, MRPS27,
CPT1A, HIBCH, APOBEC3F,
RPL17, ACTR3, VAMP1, SEC31A,
MYO6, TMEM115,ARFGEF2,
ARFGAP2, etc)
C
Group II
(Fig. 7)
GBM_Miro1-WT
HEK_Miro1-WT
DMSO
MR3
DMSO
MR3
Log2 (normalized intensity)
(ARMC10 (OMM from Mitocarta3),
GGA, TBCE, PPA1 (cytosol from org-IP),
SRRT (nucleus from org-IP),
BRCA2 (early endosome from org-IP),
NUDT1, FAM90A2P, ARL14EP,
LBR, RPS5, MCM3, etc (not in data))
D
Group IV
(Fig. 7)
GBM_Miro1-WT
HEK_Miro1-WT
DMSO
MR3
DMSO
MR3
Log2 (normalizaed intensity)
(ASCC3 (translation from org-IP),
KATNA1, SSB, CTSD, PTMA, C22orf31, NEK9 (not in data))
